# Single-Cell and Spatial Analysis Reveals Retention of FCRL4⁺ Memory B Cells in Pediatric Tonsillar Hypertrophy

**DOI:** 10.64898/2026.09.18.751603

**Authors:** Julia Young Baik, Geon Woo Park, Siyeon Jin, Jiwon Koh, Yewon Moon, Hyeewon Seo, Ha Yeon Shin, Chaeyoon Kim, Taeeun Gu, Minseok Kim, Doo Hee Han, Hyun Je Kim

**Affiliations:** Department of Biomedical Sciences, Seoul National University Graduate School, Seoul, Korea; Department of Otorhinolaryngology, Seoul National University College of Medicine, Seoul, Korea; Department of Pathology, Seoul National University Hospital, Seoul, Korea; Department of Microbiology and Immunology, Seoul National University College of Medicine, Seoul, Korea; Cancer Research Institute, Seoul National University College of Medicine, Seoul, Korea; Interdisciplinary Program in Artificial Intelligence (IPAI), Seoul National University, Seoul, Korea; School Of Transdisciplinary Innovations, Seoul National University, Seoul, Korea

**Author notes:** Corresponding author. (Doo Hee Han); Hyun Je Kim. These authors contributed equally to this work (Julia Young Baik, Geon Woo Park, Siyeon Jin). E-mail: Julia Young Baik, Geon Woo Park, Siyeon Jin, Jiwon Koh, Yewon Moon, Hyeewon Seo, Ha Yeon Shin, Chaeyoon Kim, Taeeun Gu, Minseok Kim, Doo Hee Han, Hyun Je Kim.

## Abstract

Pediatric tonsillar hypertrophy is a leading cause of sleep-disordered breathing in children, yet the immune mechanisms sustaining tonsillar enlargement remain unclear. Using integrated single-cell and spatial profiling of pediatric tonsils in a discovery cohort (n = 45) and independent validation cohort (n = 32), we identify a selective expansion and follicular retention of FCRL4⁺ memory B cells within the mantle zone of hypertrophic tonsils. These cells exhibit a quiescent, tissue-adapted phenotype characterized by NFATC1 repression and reciprocal RUNX1–RUNX2 regulation. Their accumulation is orchestrated by follicular type 1 regulatory T (Tr1) cells, which engage CTLA-4–CD86 checkpoint interactions to suppress plasma cell differentiation and reinforce a local immune tolerance program. Together, these interactions establish a persistent follicular niche enriched in regulatory FCRL4⁺ memory B cells. Our findings delineate a mechanistic Tr1–FCRL4⁺ B cell axis underlying follicular hypertrophy and highlight potential immunomodulatory targets for nonsurgical management of pediatric sleep-disordered breathing.

## Introduction

Tonsillar hypertrophy (TH) is a leading cause of pediatric upper airway obstruction, affecting up to 11% of school-aged children^1^, with over half a million tonsillectomies performed annually in the United States^2^. Tonsillar growth peaks between five and six years of age^3, 4^, coinciding with the period when adenotonsillectomy is most frequently performed^5, 6^. Hypertrophic tonsils narrow the upper airway during sleep^7^, leading to sleep-disordered breathing (SDB) that manifests as habitual snoring or mouth breathing in children^8, 9^. Chronic SDB is clinically significant, as it has been linked to neurocognitive impairment^10,11^ and reduced academic performance^12^.

Obstructive sleep apnea (OSA), the most severe manifestation of SDB, affects approximately 1–4% of children^8^ and is associated with cardiovascular complications^13, 14^. Although adenotonsillectomy remains the standard first-line treatment^15, 16^, up to 40% of patients experience residual disease^17^ and face surgical complications such as post-tonsillectomy hemorrhage^18, 19^, particularly among children under three years^20^. While systemic immune deficits after tonsillectomy are not evident^21, 22, 23^, some epidemiological studies suggest increased susceptibility to regional and long-term infections^24, 25^.

Several factors have been implicated in adenotonsillar hypertrophy and pediatric OSA, including obesity^26^, passive smoke exposure^27^, mold^28^, and microbiome alterations such as pathogenic taxa^29^ and crypt biofilms^30^. However, these associations do not fully explain the mechanisms driving tonsillar enlargement. Recently, increasing evidence suggest that immune dysregulation within the tonsillar microenvironment may represent a central pathogenic process.

Tonsils are secondary lymphoid organs (SLOs) within the mucosa-associated lymphoid tissue (MALT) system^31^, maintaining germinal center (GC) reactions essential for B cell maturation and memory formation^32, 33^. Recent single-cell and spatial studies have mapped diverse immune cell subsets and GC programs within human tonsils^34, 35^, implicating aberrant lymphoid remodeling in disease states. Histopathological analyses distinguish TH from recurrent tonsillitis by its prominent follicular hyperplasia^36, 37^, suggesting a distinct immunopathological basis. Pediatric TH exhibits heightened mucosal immune activation characterized by increased TLR4/TLR7 expression, IL-1β and NF-κB signaling^38^, and chronic immune stimulation by commensal bacteria^39^. Recent profiling further revealed expansion of activated B and Tfh-like CD4⁺ T cells^35^, enrichment of naïve CD27⁻CD21^hi B cells with reduced innate lymphoid cells^40^, and the emergence of IL-10–producing regulatory B cells that modulate local immune responses^41^.

Here, we applied an integrated single-cell and spatial multi-omic framework to reconstruct the immune architecture of pediatric tonsils across the hypertrophy spectrum. This approach delineates spatially organized immune circuits that drive lymphoid remodeling, providing mechanistic insight into TH pathogenesis and potential targets for non-surgical intervention.

## Results

### Follicular B cell expansion constitutes the architectural hallmark of hypertrophic tonsils

We stratified 77 pediatric patients by tonsil size using the Brodsky grading system, a well-validated tool for assessing TH in children^42, 43^. Clinical confounders, including body mass index (BMI), recurrent tonsillitis, SDB, upper respiratory infection (URI), and allergic rhinitis, were accounted for. Patients with an average Brodsky grade <3 were classified as having small tonsils (n = 40), and those with grade ≥ 3 as having big tonsils (n = 37). Baseline demographic and clinical characteristics were comparable between groups in both the discovery (n = 45) and validation (n = 32) cohorts (Table 1).

**Table 1.** Clinical characteristics of pediatric participants in the Discovery and Validation cohorts stratified by tonsil size. Statistical comparisons were performed using the Mann–Whitney test. No significant differences were observed.

|  | Discovery Cohort (n=45) |  |  | Validation Cohort (n=32) |  |  |
| --- | --- | --- | --- | --- | --- | --- |
|  | Small<br>(n=23) | Big<br>(n=22) | p value | Small<br>(n=17) | Big<br>(n=15) | p value |
| **Sex** |  |  | 1.000 |  |  | 1.000 |
| Male | 13 | 14 |  | 12 | 11 |  |
| Female | 10 | 8 |  | 5 | 4 |  |
| **Age (mean, range)** | 6.35 (4-13) | 6.23 (4-12) | 0.9962 | 7.63 (4-17) | 7.39 (3-10) | 0.4328 |
| **BMI (mean)** | 16.91 | 17.19 | 0.8439 | 18.95 | 18.61 | 0.4554 |
| **Tonsillitis** |  |  | 1.000 |  |  | 1.000 |
| positive | 19 | 16 |  | 9 | 5 |  |
| negative | 4 | 6 |  | 8 | 10 |  |
| **SDB** |  |  | 1.000 |  |  | 1.000 |
| positive | 14 | 20 |  | 13 | 12 |  |
| negative | 9 | 2 |  | 4 | 3 |  |
| **URI / year (mean)** | 7.96 | 8.77 | 0.4986 | 6.75 | 6.93 | 0.3505 |
| **Allergic rhinitis** |  |  | 1.000 |  |  | 1.000 |
| positive | 10 | 13 |  | 9 | 11 |  |
| negative | 13 | 9 |  | 8 | 4 |  |

To define immune correlates of tonsil size, we applied a tiered discovery–validation framework. The Discovery-Main cohort (n=16) underwent comprehensive multi-platform profiling, including single-cell RNA sequencing (scRNA-seq), GeoMx Digital Spatial Profiling (DSP), PhenoCycler multiplex imaging, immunohistochemistry (IHC) for FCRL4, and in a subset of samples, flow cytometry and CD20 IHC. The Discovery-Extended cohort (n=29) was selectively analyzed by CD20 IHC, flow cytometry, or MACSima spatial proteomics, whereas the Validation cohort (n = 32) was analyzed exclusively by scRNA-seq (Fig. 1A; Supplementary Table 1)

**Figure 1.**
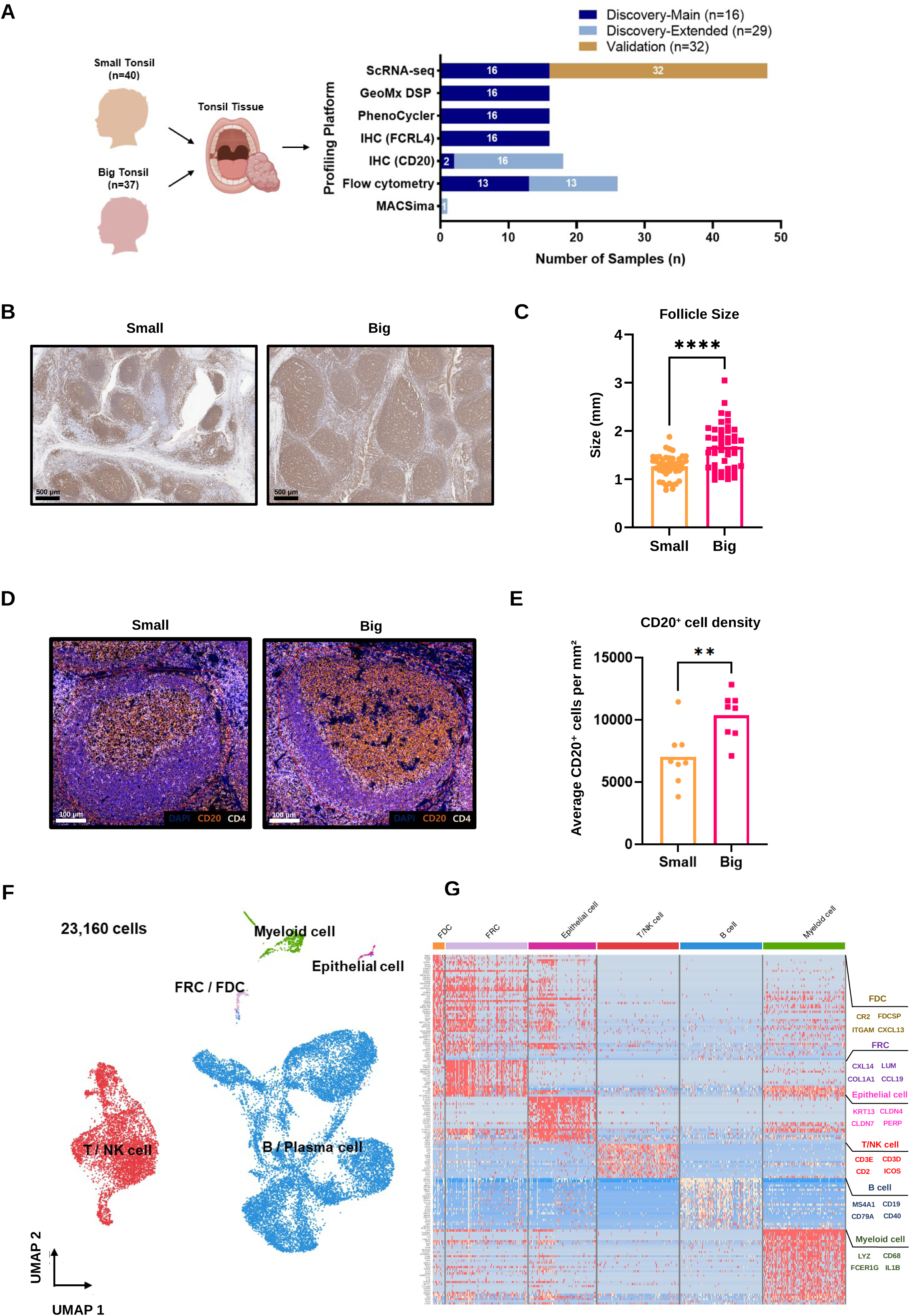
Follicular hyperplasia and B cell accumulation define hypertrophic pediatric tonsils. (A) Overview of the pediatric tonsil cohorts and the profiling platforms applied across all samples (n = 77). The bar plot indicates the number of samples analyzed by each assay within the Discovery-Main (n = 16), Discovery-Extended (n = 29), and Validation (n = 32) cohorts. (B) Representative CD20 immunohistochemistry of pediatric tonsils (scale bars, 500 μm). (C) Quantification of follicle size, defined as the long axis (mm) of individual follicles in small (n = 9) and big (n = 9) tonsils. Each dot represents one follicle (five follicles per patient). Bars indicate mean ± SD. (D) Multiplex immunofluorescence of tonsillar follicles stained with DAPI (nuclei), CD20 (B cells), and CD4 (T cells) (scale bars, 100 μm). (E) Quantification of follicular CD20⁺ cell density, calculated as the average number of CD20⁺ cells per mmZ across multiple follicles in the Discovery-Main cohort (small, n = 8; big, n = 8). Each dot represents one patient. Bars indicate mean ± SD. (F) UMAP of 23,160 single cells from the Discovery-Main cohort (n = 16), colored by annotated immune lineages. (G) Heatmap of canonical marker genes used to define the immune subsets shown in (F). Statistical significance was assessed using Mann–Whitney test in (C) and (E) (****p < 0.0001; **p < 0.01). UMAP, Uniform Manifold Approximation and Projection.

Because TH is characterized by enlargement of B-cell–rich follicles, we first examined the follicular domain. CD20, a widely used marker of mature B cells expressed from the pre-B to memory stages and downregulated upon plasma cell differentiation^44^, was used to delineate tonsillar architecture. Histologic analysis revealed prominent follicular hyperplasia with GC enlargement, constituting the architectural hallmark of hypertrophic tonsils^38, 39^, as big tonsils exhibited markedly larger follicles compared with small tonsils (p < 0.0001) (Fig. 1B, C). Multiplex immunofluorescence (mIF) provided higher-resolution spatial mapping, confirming CD20⁺ B cell localization within follicles and GCs (Supplementary Fig. 1A; Fig. 1D). Quantification of regions of interest (ROIs) encompassing follicular and limited perifollicular areas showed significantly higher CD20⁺ B cell density in big tonsils (p < 0.01) (Fig. 1E), supporting B cell accumulation as the principal histological correlate of TH.

To extend these architectural observations, we profiled the tonsillar cellular landscape by scRNA-seq. Analysis of the Discovery-Main cohort (n = 16; small, n = 8; big, n = 8) identified six major populations: B/plasma cells, T/NK cells, myeloid cells, follicular reticular cells (FRCs), follicular dendritic cells (FDCs), and epithelial cells (Fig. 1F, G). Among these, B and T/NK cells predominated, with B cells showing the highest tissue enrichment (Supplementary Fig. 1B), consistent with prior tonsillar transcriptomic atlases^34, 35^. Mirroring the pronounced follicular expansion observed by histology and multiplex imaging, scRNA-seq demonstrated a B-cell–skewed cellular landscape in hypertrophic tonsils. Although not statistically significant, B cell frequency exhibited a positive trend with tonsil size (r = 0.43, p = 0.1) (Supplementary Fig. 1C), suggesting a potential association between B cell enrichment and hypertrophic enlargement.

### Hypertrophic tonsils show selective expansion of FCRL4^+^ Memory B cells

To dissect B cell contributions to TH, we subclustered scRNA-seq data from the Discovery-Main cohort and identified 14 transcriptionally distinct subsets (Fig. 2A, B). Consistent with established models of tonsillar B-cell maturation, our BCR repertoire analysis revealed that naïve B cells predominantly carried unmutated, unswitched isotypes, whereas both class-switch recombination (CSR) and somatic hypermutation (SHM) progressively increased along the differentiation trajectory^45, 46^ (Supplementary Fig. 2A, B). Among these, proliferating FCRL4⁺ memory B cells (MBCs) were selectively expanded in hypertrophic tonsils (p < 0.05), and their frequency correlated positively with tonsil size (r = 0.58, p = 0.018), a pattern not observed in other subsets (Supplementary Fig. 2C, D). Together, the proportion of FCRL4⁺ MBCs within the memory compartment was significantly higher in big than in small tonsils (p < 0.05) and correlated with tonsil size (r = 0.51, p = 0.045) (Fig. 2C, D).

**Figure 2.**
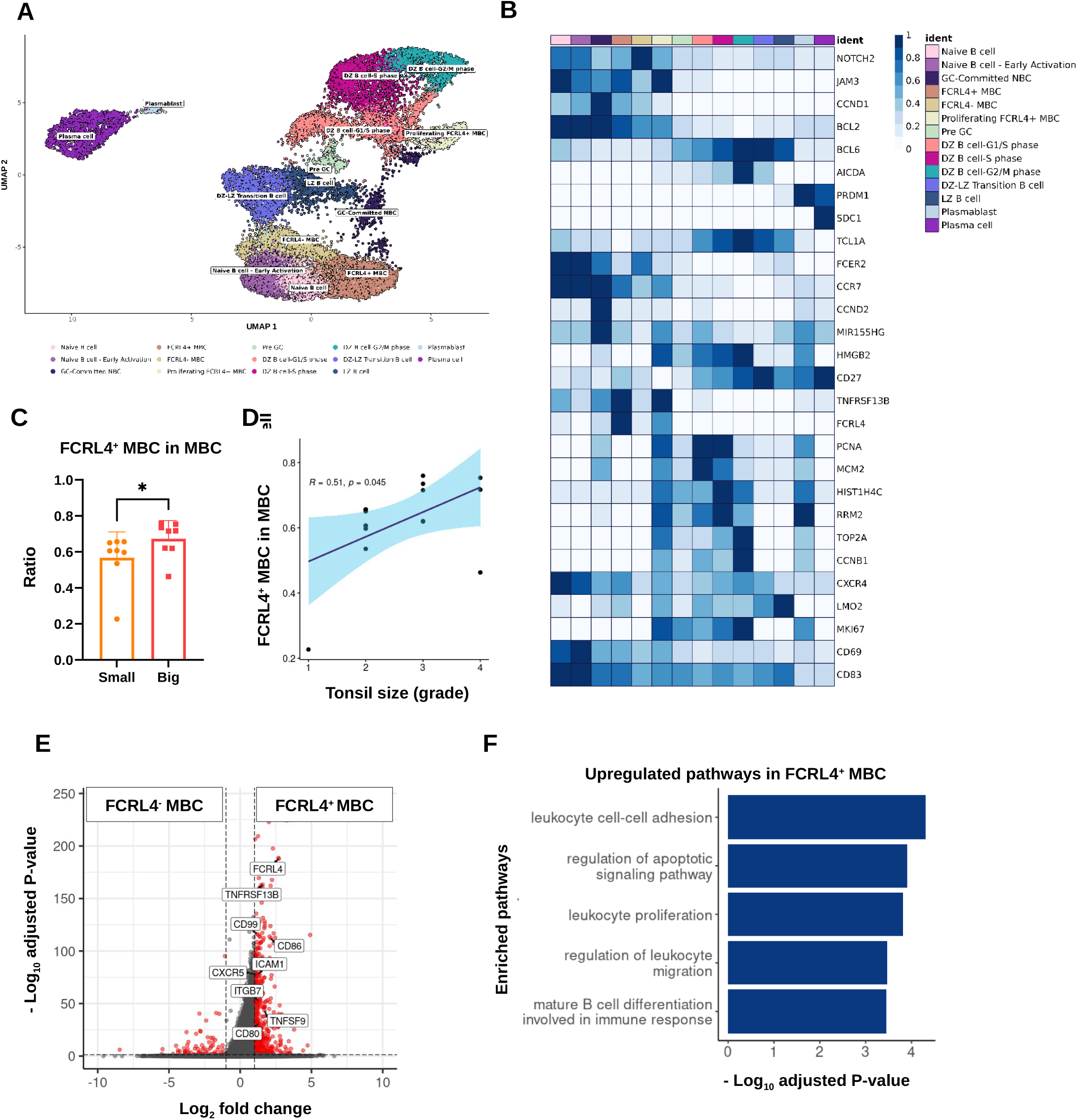
Expansion and transcriptional features of FCRL4⁺ MBCs in hypertrophic tonsils. (A) UMAP of tonsillar B cells (16,132 cells from the Discovery-Main cohort, n = 16), colored by annotated subsets. (B) Heatmap of canonical marker expression across the B cell subsets shown in (A). (C) Proportion of FCRL4⁺ MBCs among total MBCs in tonsils from the Discovery-Main cohort (small, n = 8; big, n = 8). Each dot represents one patient. Bars indicate mean ± SD. (D) Correlation between pre-operative right tonsil size and the frequency of FCRL4⁺ MBCs among total MBCs in the Discovery-Main cohort (small, n = 8; big, n = 8). Each dot represents one patient. (E) Volcano plot of DEGs between FCRL4⁺ and FCRL4⁻ MBCs, with genes upregulated in FCRL4⁺ MBCs highlighted. (F) GO enrichment analysis of genes upregulated in FCRL4⁺ MBCs. Statistical significance was assessed using Mann– Whitney test in (C) (*p < 0.05) and Spearman correlation in (D). MBC, memory B cell; DEG, differentially expressed gene; GO, Gene Ontology.

FCRL4⁺ MBCs have been described as a tissue-adapted subset in human tonsils with attenuated BCR signaling, atypical memory features, and a hyporesponsive phenotype^47, 48, 49, 50^. To characterize the transcriptional features distinguishing FCRL4⁺ from FCRL4⁻ MBCs, we analyzed differential gene expression and pathway enrichment (Fig. 2E, F). The upregulated gene set included CXCR5, ICAM1, ITGB7, and CD99, all key mediators of leukocyte migration and tissue retention, aligning with the adhesion and retention signatures previously reported in FCRL4⁺ MBCs^48, 50^. Upregulation of TNFRSF13B (TACI), a marker of epithelium-associated FCRL4⁺ B cells^51^, together with increased CD80, CD86, and TNFSF9, key co-stimulatory molecules that enhance T–B interactions and promote lymphocyte survival^52, 53^, also implicate enhanced survival and co-stimulatory signaling in this cell population. Collectively, these transcriptional changes reflect enrichment of adhesion and survival pathways, supporting the persistence of tissue-adapted FCRL4⁺ MBCs in hypertrophic tonsils.

Within this framework, FCRL4⁺ MBCs displayed a striking deviation from canonical maturation. Proliferating FCRL4⁺ MBCs remained largely unswitched with minimal SHM, resembling naïve-like states, whereas non-proliferating FCRL4⁺ MBCs acquired only partial class switching and intermediate SHM levels. In contrast, FCRL4⁻ MBCs exhibited more advanced isotype switching and markedly higher SHM frequencies, reflecting their more extensive GC experience (Supplementary Fig. 2A, B). This restricted maturation program is consistent with in vitro differentiation assays, where FCRL4⁺ B cells undergo class switching to IgG/IgA approximately two-fold less frequently than FCRL4⁻ cells^49^, and repertoire studies showing that FCRL4⁺ MBCs harbor fewer somatic mutations in their variable regions^54^. Extending these observations to TH, we found that FCRL4⁺ MBCs from hypertrophic tonsils exhibited reduced IgA⁺ and increased IgD⁺ frequencies (p < 0.05), along with a markedly lower SHM burden compared with those from small tonsils (p < 0.001) (Supplementary Fig. 2E, F), indicating constrained GC activity and incomplete maturation. Collectively, these data indicate that FCRL4⁺ MBCs remain incompletely matured and accumulate preferentially in hypertrophic tonsils.

### FCRL4⁺ Memory B Cells accumulate within follicular niches of hypertrophic tonsils

To delineate the spatial distribution of FCRL4⁺ MBCs, we performed immunohistochemistry on Discovery-Main tonsils. We annotated GC and mantle zone boundaries in five follicles per sample to focus on follicular regions, as hypertrophic tonsils displayed marked follicular enlargement in the preceding figures (Supplementary Fig. 3). FCRL4⁺ cells were observed both within and outside follicles ^47, 55^ (Fig. 3A).

**Figure 3.**
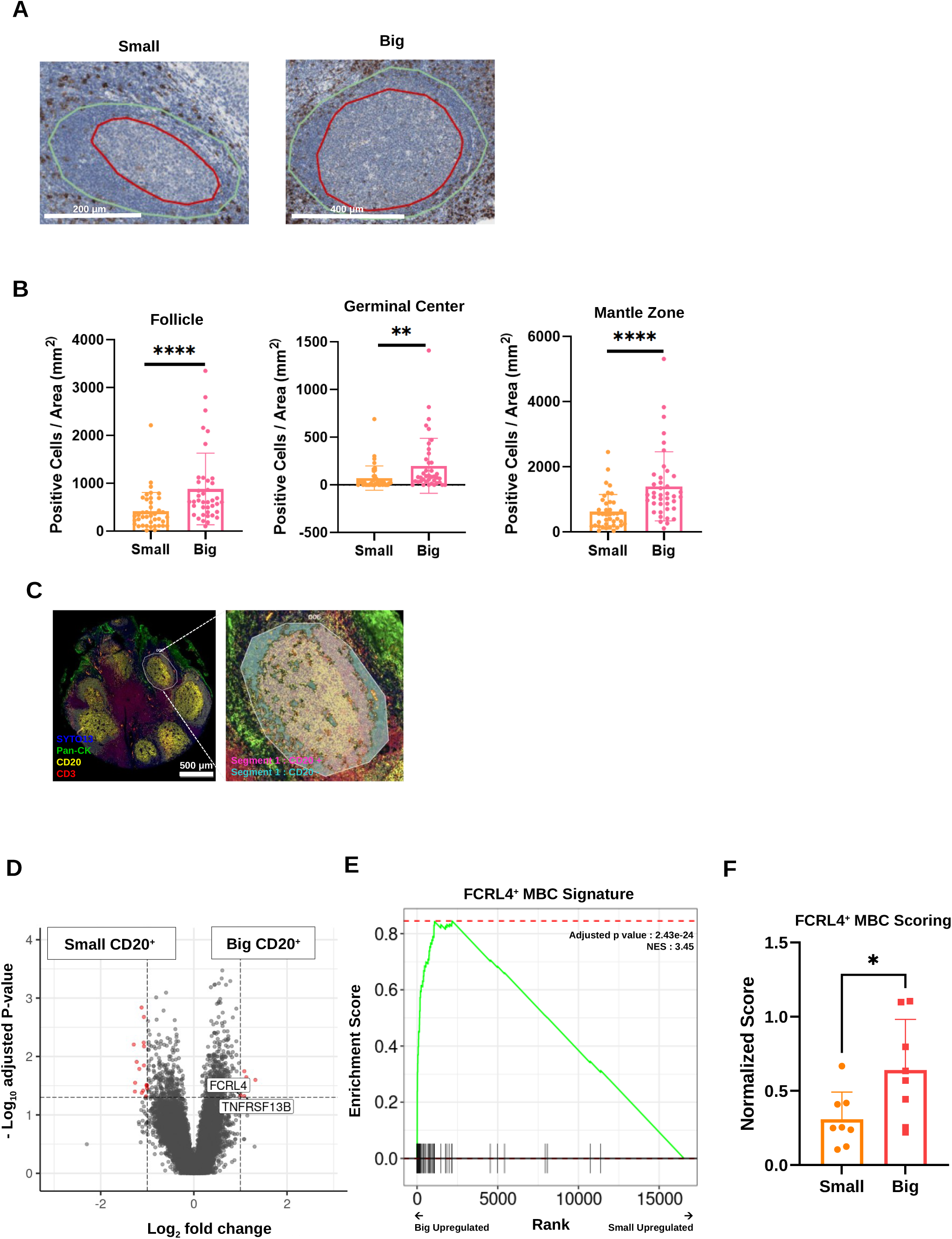
Spatial enrichment of FCRL4⁺ MBCs in hypertrophic tonsils revealed by immunohistochemistry and GeoMx profiling. (A) Representative immunohistochemistry images of FCRL4-stained follicles from small and big tonsils. Follicular boundaries were annotated to delineate the germinal center (red) and mantle zone (green) (scale bars, 200 μm). (B) Quantification of FCRL4⁺ cell density in the follicle, germinal center, and mantle zone of tonsils from the Discovery-Main cohort (small, n = 8; big, n = 8; five follicles per patient). Each dot represents one follicle. Bars indicate mean ± SD. (C) Representative GeoMx DSP images of tonsils stained with SYTO13 (nuclei), Pan-CK (epithelial cells), CD20 (B cells), and CD3 (T cells), showing follicular segmentation into CD20⁺ and CD20⁻ compartments (scale bars, 200 μm). (D) Volcano plot of DEGs between CD20⁺ segments from tonsils in the Discovery-Main cohort (small, n = 8; big, n = 8). Genes upregulated in big tonsil CD20⁺ segments are shown in red. (E) GSEA of the FCRL4⁺ MBC signature in CD20⁺ segments from tonsils in the Discovery-Main cohort (small, n = 8; big, n = 8). (F) ssGSEA scores for the FCRL4⁺ MBC signature in CD20⁺ segments from tonsils in the Discovery-Main cohort (small, n = 8; big, n = 8). Each dot represents one patient. Bars indicate mean ± SD. Statistical significance was assessed using Mann– Whitney test in (B) and (F) (*p < 0.05; **p < 0.01; ****p < 0.0001) and GSEA in (E). DSP, digital spatial profiling; GSEA, Gene Set Enrichment Analysis; ssGSEA, single-sample Gene Set Enrichment Analysis.

Hypertrophic tonsils exhibited a marked increase in FCRL4⁺ cell density across all follicular compartments, with significant enrichment in the follicle (p < 0.0001), GC (p < 0.01), and most prominently in the mantle zone (p < 0.0001) (Fig. 3B). Given that the follicle comprises both the GC and mantle zone, the overall follicular increase is likely driven predominantly by mantle zone accumulation. These findings align with our scRNA-seq data, linking FCRL4⁺ MBC expansion with preferential follicular accumulation.

To extend these findings, we applied GeoMx Digital Spatial Profiling (DSP) to tonsillar follicles (fig. S4A). ROIs encompassing follicular and interfollicular regions were segmented by CD20 expression, clearly distinguishing B cell–rich (CD20⁺) and non–B cell (CD20⁻) compartments (Fig. 3C). DEG and pathway analyses confirmed the validity of the segmentation strategy, revealing enrichment of B cell– related genes and pathways in CD20⁺ regions, whereas non–B cell transcripts predominated in CD20⁻ areas (Supplementary Fig. 4B, C). Within CD20⁺ regions, hypertrophic tonsils exhibited upregulation of FCRL4 and TNFRSF13B, both of which are upregulated genes in FCRL4⁺ MBCs according to our scRNA-seq analysis (Fig. 3D; Fig. 2E). Using a scRNA-seq–derived FCRL4⁺ MBC signature validated by gene set scoring (Supplementary Fig. 4D), we performed gene set enrichment analysis (GSEA) on DEGs generated from GeoMx DSP CD20⁺ segments. This analysis revealed significant enrichment of the FCRL4⁺ MBC program in hypertrophic follicles (NES = 3.45, adjusted p = 2.43 × 10⁻Z⁴) (Fig. 3E). Single-sample GSEA (ssGSEA) further demonstrated elevated FCRL4⁺ MBC scores in hypertrophic ROIs (p < 0.05; Fig. 3F), with a positive but nonsignificant correlation with tonsil size (r = 0.43, p = 0.1; Supplementary Fig. 4E). Together, these data show that FCRL4⁺ MBCs are both transcriptionally programmed and anatomically concentrated within hypertrophic follicles.

### FCRL4⁺ Memory B Cell expansion in hypertrophic tonsils is independent of follicular helper T cells

Given the increased abundance of FCRL4⁺ MBCs within follicles and their reduced isotype switching and SHM in the big tonsil group, we hypothesized that impaired T follicular helper (Tfh)-mediated help could underlie this phenotype (Supplementary Fig. 2E, F). This is supported by the central role of Tfh cells in B-cell maturation, germinal center formation, class-switch recombination, and affinity maturation^56, 57^.

Subclustering of T/NK cells from scRNA-seq data identified Tfh cells within tonsillar tissue (Fig. 4A, B). Cell–cell interaction analysis further demonstrated that Tfh cells showed preferential connectivity with FCRL4⁺ MBCs, suggesting close spatial and functional coupling between these two compartments (Fig. 4C). However, Tfh frequencies were comparable between small and big tonsils (p > 0.05; Fig. 4D) and showed no significant correlation with preoperative right tonsil size (r = 0.12, p = 0.66; Fig. 4E). DEG analysis detected only three differentially expressed genes, indicating minimal transcriptional divergence (Fig. 4F).

**Figure 4.**
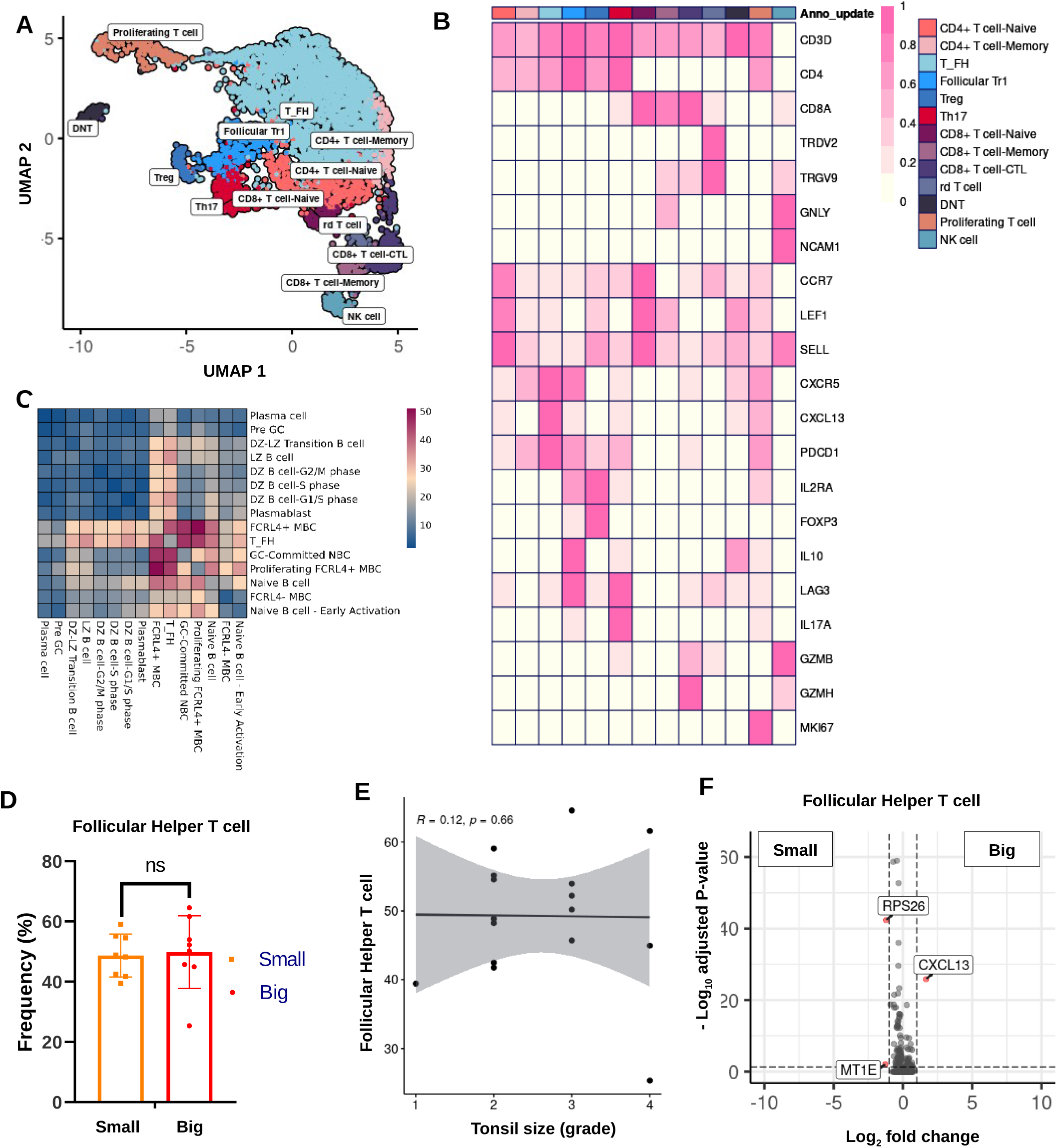
Tfh cells show no abundance or transcriptional differences by tonsil size. (A) UMAP of T/NK cell subsets (6,010 cells from the Discovery-Main cohort, n = 16), colored by annotated subsets. (B) Heatmap of canonical gene expression patterns across the T/NK subsets shown in (A). (C) Cell–cell interaction analysis between Tfh and B cell subsets in the Discovery-Main cohort (n = 16). (D) Frequency of Tfh cells in tonsils from the Discovery-Main cohort (small, n = 8; big, n = 8). Each dot represents one patient. Bars indicate mean ± SD. (E) Correlation between Tfh cell frequency and pre-operative right tonsil size in the Discovery-Main cohort (small, n = 8; big, n = 8). Each dot represents one patient. (F) Volcano plot of DEGs in Tfh cells comparing tonsils in the Discovery-Main cohort (small, n = 8; big, n = 8). Statistical significance was assessed using Mann–Whitney test in (D) and Spearman correlation in (E). No significant differences were observed. UMAP, Uniform Manifold Approximation and Projection; DEG, differentially expressed gene; Tfh, T follicular helper cell.

To assess these findings spatially, we generated a Tfh signature from scRNA-seq data (Supplementary Fig. 5A) and applied it to GeoMx DSP CD20⁻ segments, which represent non–B cell regions. This analysis revealed no group-specific differences in Tfh enrichment, as GSEA demonstrated no significant enrichment of the Tfh signature (NES = 0.98, adjusted p = 0.48; Supplementary Fig. 5B). GeoMx-based scoring also confirmed comparable Tfh levels between small and big tonsils (p > 0.05; Supplementary Fig. 5C). Flow cytometry further confirmed comparable Tfh frequencies between small and big tonsils (p > 0.05; Supplementary Fig. 5D, F). Expression of canonical functional markers ICOS, PD-1, and BCL6 was indistinguishable between groups (p > 0.05; Supplementary Fig. 5E, G), consistent with the established ICOS–BCL6 axis that defines Tfh identity^58^. Collectively, these results indicate that Tfh cells in hypertrophic tonsils are quantitatively and qualitatively preserved, suggesting that expansion of FCRL4⁺ MBCs is driven by alternative immune circuits.

### A Tr1–FCRL4⁺ Memory B Cell interaction axis in hypertrophic tonsils

To dissect Tfh-independent mechanisms of FCRL4⁺ MBC expansion, we profiled T cell subsets between groups. Among all subsets, naïve CD4⁺ T cells were significantly reduced (p < 0.01) (Supplementary Fig. 6A), whereas follicular type 1 regulatory T (Tr1) were increased (p < 0.05) in hypertrophic tonsils (Fig. 5A). Follicular Tr1 cells, defined by their IL-10–dependent immunosuppressive function and co-expression of CD49b and LAG-3, have been identified in human tonsillar tissue^59, 60^. To distinguish follicular Tr1 cells from FOXP3⁺ follicular regulatory T (Tfr) cells — classical germinal center–associated regulatory T cells that suppress B cell responses—multiplex imaging was performed. This analysis confirmed that Tfr cells were absent within follicles (Supplementary Fig. 6H), excluding canonical FOXP3+ Tfr cells as the source of the observed regulatory activity. Consistent with their selective enrichment, follicular Tr1 frequency correlated positively with pre-operative right tonsil size (Fig. 5B). Trajectory analysis indicated a progressive transition from naïve CD4⁺ T cells toward follicular Tr1 cells (Supplementary Fig. 6B), with their frequencies exhibiting a negative correlation (r = –0.41, p = 0.12; Supplementary Fig. 6C). These results suggest an increased abundance of Tr1 cells in the tonsil big group.

**Figure 5.**
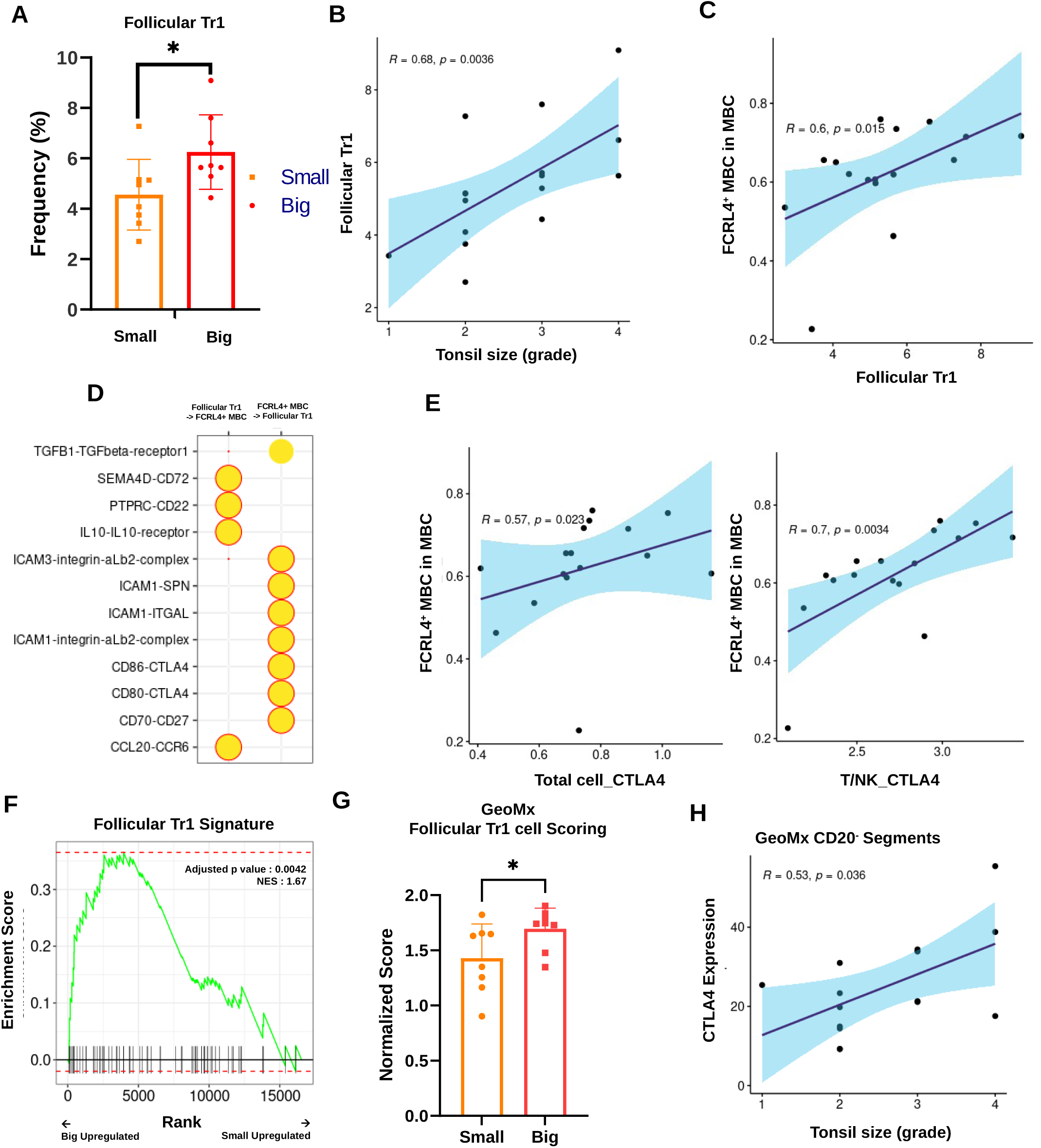
Follicular Tr1–FCRL4⁺ MBC axis expands in hypertrophic tonsils via CTLA-4–mediated regulation. (A) Frequency of follicular Tr1 cells in tonsils from the Discovery-Main cohort (small, n = 8; big, n = 8). Each dot represents one patient. Bars indicate mean ± SD. (B) Correlation between pre-operative right tonsil size and follicular Tr1 frequency in the Discovery-Main cohort (small, n = 8; big, n = 8). Each dot represents one patient. (C) Correlation between follicular Tr1 frequency and the proportion of FCRL4⁺ MBCs among memory B cells in the Discovery-Main cohort (small, n = 8; big, n = 8). Each dot represents one patient. (D) Predicted ligand–receptor interactions between follicular Tr1 cells and FCRL4⁺ MBCs from scRNA-seq data. (E) Correlation between FCRL4⁺ MBC frequency and CTLA-4 expression in total T cells (left) and T/NK cells (right) from the Discovery-Main cohort (small, n = 8; big, n = 8). Each dot represents one patient. (F) GSEA of the follicular Tr1 signature in CD20 ⁻ segments from tonsils in the Discovery-Main cohort (small, n = 8; big, n = 8). (G) ssGSEA scores for the follicular Tr1 signature in CD20⁻ segments from tonsils in the Discovery-Main cohort (small, n = 8; big, n = 8). Each dot represents one patient. Bars indicate mean ± SD. (H) Correlation between pre-operative right tonsil size and CTLA-4 expression in CD20⁻ segments from the Discovery-Main cohort (small, n = 8; big, n = 8). Each dot represents one patient. Statistical significance was assessed using Mann– Whitney test in (A) and (G) (*p < 0.05) and Spearman correlation in (B), (C), (E), and (H). Tr1, type 1 regulatory T cell; MBC, memory B cell; GSEA, Gene Set Enrichment Analysis; ssGSEA, single-sample Gene Set Enrichment Analysis.

DEG analysis of Tr1 revealed enrichment of B cell–regulatory pathways, including activation, proliferation, and differentiation, suggesting that Tr1 cells may directly influence GC B cell responses (Supplementary Fig. 6D). We next examined the relationship between Tr1 cells and FCRL4⁺ MBCs. Tr1 frequency was positively correlated with FCRL4⁺ MBC enrichment (r = 0.60, p = 0.015; Fig. 5C). Cell– cell interaction analysis revealed significant IL-10–IL10RA^59^, CTLA-4–CD86^61^, and TGFB1– TGFBR1^62^ interactions from follicular Tr1 to FCRL4⁺ MBCs B cells (Fig. 5D, Supplementary Fig. 6E). These reciprocal cytokine and costimulatory patterns suggest a potential interaction circuit between Tr1 cells and tissue-resident FCRL4⁺ MBCs within the follicular niche, which may influence FCRL4⁺ MBC behavior.

To identify Tr1-derived factors potentially linked to the expansion of FCRL4⁺ MBCs, we examined genes selectively expressed by follicular Tr1 cells and found CTLA-4 as a prominent candidate. Re-examination of our single-cell dataset confirmed that CTLA-4 expression was restricted to the T/NK lineage and peaked in the follicular Tr1 subset (Supplementary Fig. 6F). Moreover, CTLA-4 levels were positively correlated with FCRL4⁺ MBC frequency (total CTLA-4: r = 0.57, p = 0.023; T/NK CTLA-4: r = 0.70, p = 0.0034; Fig. 5E). Among the CTLA-4 ligands, CD86 was selectively upregulated in FCRL4⁺ MBCs (Supplementary Fig. 6G).

Given the increased abundance of follicular Tr1 cells by scRNA-seq, we evaluated their spatial enrichment using GeoMx DSP. GSEA confirmed significant enrichment of the follicular Tr1 signature in hypertrophic tonsils, specifically within CD20⁻ segments (NES = 1.67, adjusted p = 0.0042; Supplementary Fig. 6I, Fig. 5F). Consistently, Tr1 signature scores were elevated across hypertrophic ROIs (p < 0.05; Fig. 5G). Spatial transcriptomics additionally revealed that CTLA-4 expression in CD20⁻ regions positively correlated with preoperative right tonsil size (r = 0.53, p = 0.036; Fig. 5H).

To directly assess the spatial relationship between follicular Tr1 cells and FCRL4⁺ MBCs, we generated Xenium spatial transcriptomic profiles from a tonsillar tissue microarray comprising 15 samples (Supplementary Fig. 7A). Cell-type annotation based on canonical marker expression resolved 23 cell populations, including follicular Tr1 cells and FCRL4⁺ MBCs, with transcriptional profiles consistent with those identified by scRNA-seq (Supplementary Fig. 7B, C). Spatial visualization revealed frequent close localization of follicular Tr1 cells and FCRL4⁺ MBCs within tonsillar follicles (Supplementary Fig. 7D). Consistent with this observation, neighborhood enrichment analysis showed that follicular Tr1 cells were preferentially associated with FCRL4⁺ MBCs compared with FCRL4⁻ MBCs (p = 0.0043; Supplementary Fig. 7D). Together, these findings indicate that follicular Tr1 cells represent a prominent interacting partner of FCRL4⁺ MBCs and may contribute to shaping the immune environment of hypertrophic tonsils.

### Follicular Tr1 cells impede FCRL4⁺ Memory B Cell differentiation

Given the increased abundance of follicular Tr1 cells in big tonsils (Fig. 5), we next investigated how exactly these cells influence FCRL4⁺ MBCs. Trajectory analysis of the tonsillar B cell compartment revealed a branch in which FCRL4⁺ MBCs transition through an FCRL4⁻ intermediate (Fig. 6A), consistent with prior reports demonstrating that the FCRL4⁺ state is a reversible, activation-dependent phenotype that can remodel back to an FCRL4⁻ memory state^48, 49^. Consistent with impaired effector differentiation FCRL4⁺ MBC abundance showed an inverse correlation with plasma cells (r = –0.52, p = 0.04) (Fig. 6B), supporting impaired differentiation toward effector fates in hypertrophic tonsils.

**Figure 6.**
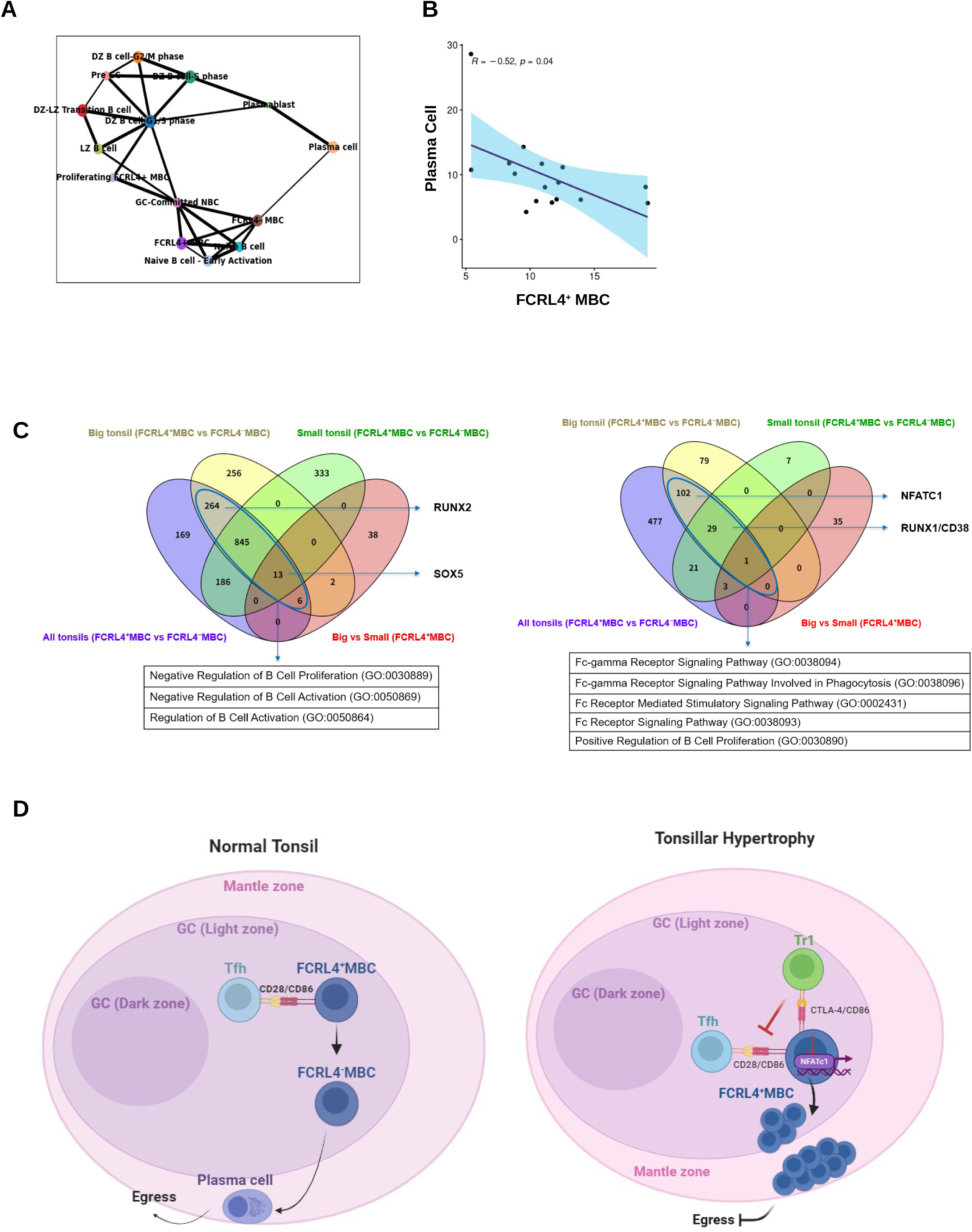
Follicular Tr1–FCRL4⁺ MBC axis restrains NFATC1 signaling and plasma cell differentiation in hypertrophic tonsils. (A) Trajectory analysis of tonsillar B cells from the Discovery-Main cohort (n = 16), showing canonical differentiation from naive and germinal center states toward plasmablasts and plasma cells, with an alternative branch leading to FCRL4⁺ MBCs. (B) Correlations between the frequency of FCRL4⁺ MBCs and plasma cells in the Discovery-Main cohort (small, n = 8; big, n = 8). Each dot represents one patient. (C) Venn diagrams of differentially expressed genes (DEGs) in FCRL4⁺ MBCs from big versus small tonsils in the Discovery-Main cohort (small, n = 8; big, n = 8), with Gene Ontology enrichment analysis of upregulated and downregulated gene sets. (D) Schematic model illustrating how Tr1-derived CTLA-4 suppresses NFATC1 in FCRL4⁺ MBCs via CD86, leading to their accumulation. Statistical significance was assessed using Spearman correlation in (B). Tr1, type 1 regulatory T cell; MBC, memory B cell; DEG, differentially expressed gene.

To define transcriptional features associated with tonsillar hypertrophy, we compared FCRL4⁺ and FCRL4⁻ memory B-cell subsets across big and small tonsils using four complementary DEG analyses. These included FCRL4⁺ vs FCRL4⁻ (pooled), Big FCRL4⁺ vs Big FCRL4⁻, Small FCRL4⁺ vs Small FCRL4⁻, and Big FCRL4⁺ vs Small FCRL4⁺, which were integrated by Venn analysis to identify conserved and hypertrophy-specific signatures (Fig. 6C; Supplementary Table 2, 3). Integration of the four DEG comparisons revealed a set of genes that were consistently upregulated in FCRL4⁺ MBCs, including those showing higher expression specifically in big tonsils. These upregulated genes were enriched for pathways related to the negative regulation of B cell activation and proliferation, indicating a restrained activation program characteristic of tissue adaptation. Among the transcription factors enriched in hypertrophic tonsils, RUNX2 emerged as a key regulator reinforcing the tissue-adapted phenotype of FCRL4⁺ MBCs. RUNX2 expression was selectively elevated in FCRL4⁺ MBCs from big tonsils, consistent with prior work identifying the SOX5–RUNX2 axis as a regulator of quiescent, tissue-adapted memory B cells^48, 63^. In contrast, SOX5 was robustly expressed in FCRL4⁺ MBCs from both small and big tonsils, marking a conserved core program independent of tonsil size.

Conversely, genes that were consistently downregulated in FCRL4⁺ MBCs across the integrated DEG analyses were enriched for pathways related to Fc receptor signaling and B-cell proliferation, reflecting a dampened activation state. RUNX1, a transcription factor associated with activation-associated memory B-cell differentiation, together with the activation marker CD38, was uniformly reduced in FCRL4⁺ MBCs^48, 64^. Notably, NFATC1, a transcription factor downstream of CD86 signaling and indispensable for B cell activation and plasmablast commitment^65^, was selectively downregulated in FCRL4⁺ MBCs from big tonsils. We highlight NFATC1 as central because its loss, together with reciprocal RUNX1–RUNX2 regulation, likely shifts B cell fate from plasmablast differentiation toward a tissue-retained FCRL4⁺ program. In T cells, NFAT is known to directly bind and transactivate the RUNX1 promoter through a calcineurin-dependent pathway^66^ and form transcriptional complexes with RUNX1 and other nuclear partners^67^. These prior studies imply that a similar NFAT–RUNX regulatory mechanism may be conserved in B cells.

Together, these findings support a model in which FCRL4⁺ MBCs normally progress through FCRL4⁻ intermediates toward activated effector states, but in hypertrophic tonsils this transition is specifically curtailed by CTLA-4–CD86–mediated inhibition. Follicular Tr1 cells expressing CTLA-4 engage CD86 on FCRL4⁺ MBCs, a mechanism that allows CTLA-4 to outcompete CD28 and remove CD86 via transendocytosis, thereby depriving B cells of the co-stimulatory input required for NFATC1 activation. Loss of this pathway suppresses NFATC1-dependent differentiation cues and promotes retention of FCRL4⁺ MBCs within the mantle zone (Fig. 6D).

### Independent Validation Confirms FCRL4⁺ Memory B Cell Enrichment in Hypertrophic Tonsils

To validate the Discovery-cohort findings, we analyzed an independent scRNA-seq dataset of 32 tonsil samples (Table 1). Cell type annotation was performed using the SingleR algorithm, referencing the Discovery dataset for label transfer (Fig. 7A, Supplementary Fig. 8A). A total of 42,113 cells were annotated, and cross-cohort transcriptomic similarity confirmed the robustness and reproducibility of this approach (Fig. 7B, C).

**Figure 7.**
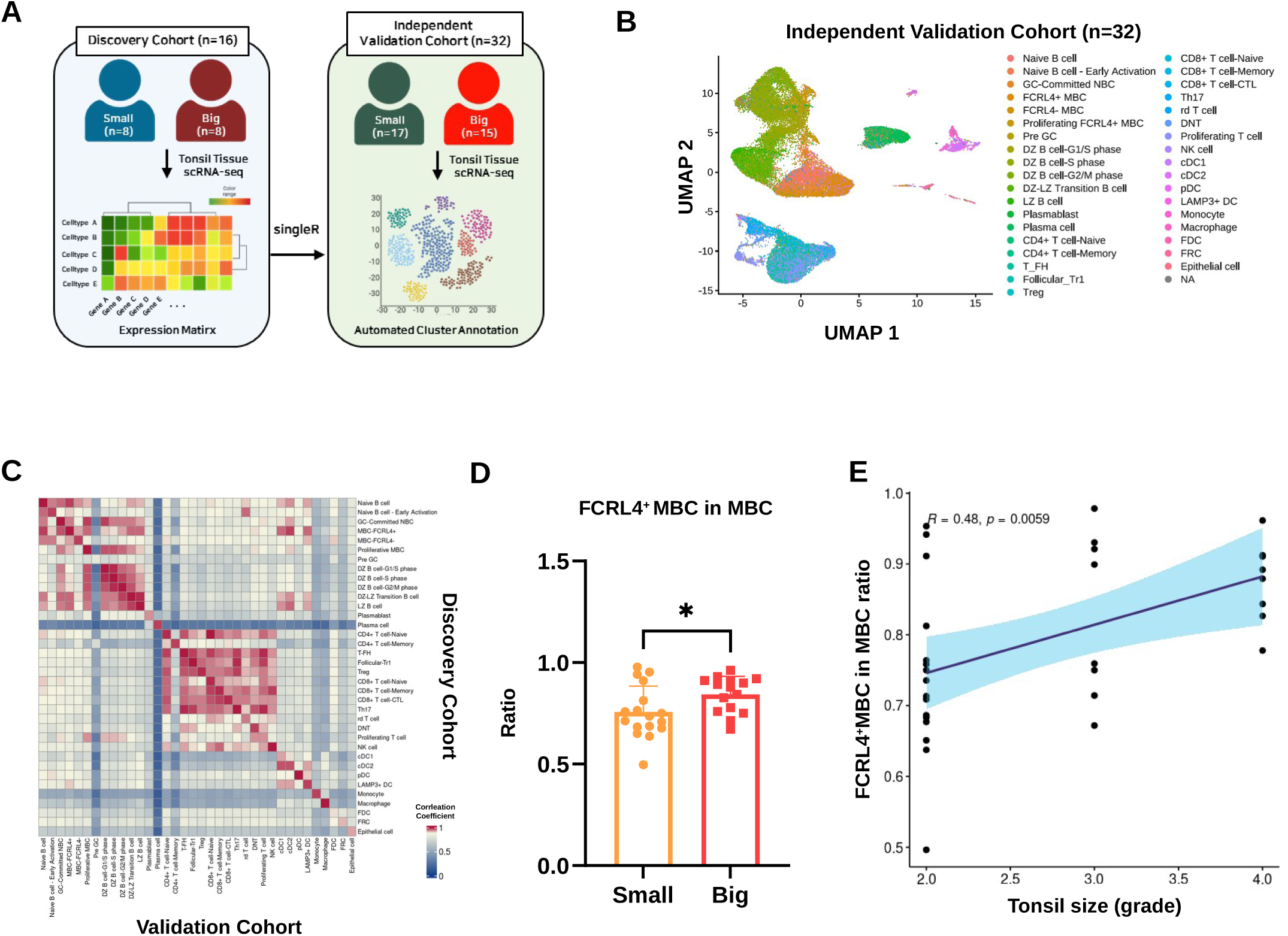
Independent cohort validation confirms enrichment of FCRL4⁺ MBCs in hypertrophic tonsils. (A) Schematic of the Discovery-Main (n = 16) and independent Validation (n = 32) cohorts for scRNA-seq. Cell type annotation in the Validation cohort was performed using SingleR with the Discovery-Main dataset as reference. (B) UMAP of 42,113 cells from the Validation cohort annotated into immune cell subsets. (C) Correlation matrix of cluster identities between the Discovery-Main (n = 16) and Validation (n = 32) cohorts. (D) Proportion of FCRL4⁺ MBCs among total MBCs in tonsils from the Validation cohort (small, n = 17; big, n = 15). Each dot represents one patient. Bars indicate mean ± SD. (E) Correlation between the proportion of FCRL4⁺ MBCs and pre-operative right tonsil size in the Validation cohort (small, n = 17; big, n = 15). Each dot represents one patient. Statistical significance was assessed using Mann–Whitney test in (D) (*p < 0.05) and Spearman correlation in (E). UMAP, Uniform Manifold Approximation and Projection; MBC, memory B cell.

FCRL4⁺ MBCs were significantly enriched in hypertrophic tonsils within the Validation cohort (p < 0.05), recapitulating the Discovery cohort dataset (Fig. 7D). Their frequency also correlated positively with pre-operative right tonsil size (r = 0.48, p = 0.0059; Fig. 7E). Follicular Tr1 cells exhibited a similar directional pattern: although their enrichment did not reach statistical significance (p = 0.14), they trended higher in hypertrophic tonsils and showed a positive, though nonsignificant, correlation with preoperative right tonsil size (r = 0.30, p = 0.098; Supplementary Fig. 8B, C). Together, these findings demonstrate that the principal cellular signatures identified in the Discovery cohort are reproducible in an independent dataset.

Building on this cross-cohort concordance, integrative modeling highlighted a regulatory circuit in which CTLA-4 expressed by follicular Tr1 cells engages CD86 on FCRL4⁺ MBCs, leading to NFATC1 suppression and impaired plasma cell differentiation. This interaction promotes the preferential retention of FCRL4⁺ MBCs within the mantle zone, establishing a tolerance-oriented follicular niche. Collectively, the multi-cohort and multi-platform analyses support a unified mechanistic model in which FCRL4⁺ MBCs act as central effectors of pediatric tonsillar hypertrophy, with follicular Tr1 cells functioning as upstream regulators that maintain these cells within the follicular niche.

## Discussion

Although TH is a common cause of adenotonsillectomy in children, the immunological processes that sustain this condition remain poorly defined. Although the initial trigger that causes tonsillar enlargement remains uncertain, our findings indicate that once hypertrophy develops, a localized immunoregulatory circuit actively sustains and amplifies follicular growth rather than resolving it. Specifically, we identify a tolerance-skewed follicular remodeling program orchestrated by the FCRL4 MBC–follicular Tr1 axis, which suppresses productive GC differentiation and promotes the persistence of enlarged follicles. This positions TH not as a purely inflammatory disorder but as a self-reinforcing immunoregulatory state, wherein a dysregulated balance between activation and tolerance drives chronic follicular persistence.

Our data reveal a CTLA-4–dependent inhibitory checkpoint linking Tr1–FCRL4⁺ MBC interaction that engages an NFATC1-driven regulatory circuit. Follicular Tr1 cells expressing CTLA-4 engage CD86⁺ FCRL4⁺ MBCs, directly competing with Tfh CD28 co-stimulation. Because CTLA-4 binds to CD86 with approximately ten-fold higher affinity than to CD28 ^52, 68^, Tr1 cells gain a clear competitive advantage to Tfh cells at the T–B interface. Furthermore, CTLA-4 not only blocks co-stimulation but also actively removes CD86 from the B-cell surface through transendocytosis, thereby sustaining prolonged costimulatory deprivation^52^. This inhibitory contact suppresses NFATC1^65^, preventing plasmablast differentiation and stabilizing the FCRL4⁺ state within hypertrophic follicles. This sustained inhibitory axis uncouples follicular expansion from productive immune activation, a defining feature of pediatric TH.

FCRL4⁺ MBCs were originally described as tissue-localized, hyporesponsive memory cells with attenuated BCR and CD40 signaling^47, 48^. Subsequent studies revealed their pro-inflammatory potential during chronic infection^69^ and ultimately redefined them as tissue-adapted memory cells rather than exhausted ones^50^. FCRL4⁺ MBCs expand across diverse pathological contexts —including HIV infection^69^, rheumatoid arthritis^70^, and Sjögren’s syndrome^51, 71^ —highlighting their propensity to accumulate at inflamed or epithelial interfaces under chronic immune activation. Mechanistically, FCRL4⁺ MBCs attenuate BCR signaling by engaging ITIM-mediated recruitment of SHP-1/2 phosphatases^47, 72^ and bind IgA^73, 74^. Within the human tonsils, FCRL4⁺ MBCs, previously identified as IRTA1⁺ B cells, reside in intraepithelial and subepithelial marginal zone niches^55^ and, in our data, show marked enrichment in the follicular mantle zone.

At the transcriptional level, FCRL4⁺ MBCs are shaped by SOX5 and RUNX2^48^, which promote long-lived, low-responsive programs that enhance cell survival under chronic stimulation while restricting plasmablast differentiation. In parallel, suppression of NFATC1, a key integrator of co-stimulatory signaling required for B cell proliferation and plasma cell commitment, provides a mechanistic basis for this restrained activation state. In hypertrophic tonsils, the combined downregulation of NFATC1, loss of RUNX1^48, 64^ and induction of RUNX2^48^, collectively shift FCRL4⁺ MBCs away from effector differentiation toward a tissue-retained, regulatory phenotype. Together, these features define FCRL4⁺ MBCs as tissue-adapted regulatory sentinels that stabilize a hyporesponsive follicular ecosystem and, in hypertrophic tonsils, contribute to the maintenance of an enlarged yet functionally quiescent architecture.

Importantly, this phenotype extends beyond the tonsil and reflects a conserved organizational principle of mucosa-associated lymphoid tissues (MALT). FCRL4⁺ MBCs are consistently observed in MALT sites such as Peyer’s patches^55^, and analogous tissue-retained MBC populations have also been described in nasal polyps^75^, where chronic stimulation similarly constrains B-cell maturation. This cross-tissue recurrence indicates that the restrained, regulatory remodeling we observe in pediatric TH is not tonsil-specific but exemplifies a broader mucosal tolerance program.

Our analyses likewise identify follicular Tr1 cells as non-canonical regulators within the germinal microenvironment. While Tfh cells showed no quantitative or transcriptional expansion in hypertrophic tonsils, follicular Tr1 cells were enriched and expressed IL-10, TGFBR1, and CTLA-4, whose corresponding receptors were detected on FCRL4⁺ MBCs. These cytokines and checkpoint signals collectively promote B-cell survival while restraining differentiation^61, 62, 76^. Tr1 cells, defined by CD49b/LAG-3 co-expression and IL-27 induction^59, 77, 78^, have been reported to expand in chronic inflammation^79^ and tumors^80^. In hypertrophic tonsils, Tr1 cells compete with Tfh cells for access to FCRL4⁺ MBCs via CTLA-4–CD86 engagement, an interaction presumed to occur predominantly within the mantle zone where FCRL4⁺ B cells preferentially accumulate.

Prior pediatric TH studies emphasized bulk-tissue proinflammatory signatures (IL-6, IL-8, TNF) ^38, 39^ and Tfh-like CD4⁺ expansion with naïve B-cell enrichment^40^, shaping a hyperreactive interpretation. By stratifying samples according to tonsil size and applying integrated single-cell and spatial profiling, our study provides a higher-resolution view of the immune architecture. Instead of diffuse inflammation, we reveal a spatially compartmentalized tolerance program within follicles, characterized not by quantitative or transcriptional expansion of Tfh cells but by selective enrichment of Tr1 cells and accumulation of FCRL4⁺ MBCs with reduced NFATC1 activity. Mechanistically, these features delineate a tolerance-dominant follicular circuit in which Tr1-derived inhibitory signals intersect with attenuated BCR/CD40–NFATC1 activity in FCRL4⁺ B cells. Together, this framework reconciles bulk inflammatory readouts with a follicle-localized inhibitory circuit, explaining why follicles remain large yet immunologically quiescent in pediatric TH.

Our conclusions are supported by a rigorous multi-platform design integrating single-cell, spatial, and proteomic analyses with independent cohort validation to ensure robustness and reproducibility. By anchoring transcriptional programs within their spatial and anatomical context, we directly link cellular states to tissue-level pathology, establishing a mechanistic framework with translational relevance. These findings also suggest a potential immunotherapeutic axis—the CTLA-4–CD86–NFATC1 pathway—as a modifiable inhibitory checkpoint in chronic mucosal hypertrophy. Targeting CTLA-4 or restoring NFATC1 activity in situ may attenuate follicular overgrowth without ablating tonsillar immune function.

While this cross-sectional study delineates a coherent model of tolerance-based follicular remodeling, it remains observational and therefore cannot establish causality. Future studies should include functional validation using ex vivo tonsil slices or organoid cultures that preserve native architecture to test whether CTLA-4–CD86 interactions directly restrain NFATC1 activation and plasma cell differentiation. Perturbation experiments—such as blocking CTLA-4 or pharmacologically restoring NFATC1 activity—could also determine whether disrupting this axis reverses the hyporesponsive phenotype of FCRL4⁺ MBCs. In parallel, co-culture assays with sorted Tr1 and FCRL4⁺ MBC subsets would enable quantitative assessment of cytokine-and checkpoint-mediated regulation at the single-cell level. Comparative analyses across pediatric cohorts, spanning recurrent tonsillitis and hypertrophic tonsils may further elucidate how this regulatory circuit evolves under chronic stimulation. Finally, integrating spatial proteomics with imaging mass spectrometry can resolve the cytokine and metabolic gradients that reinforce Tr1–FCRL4⁺ interactions in situ.

Despite these limitations, our findings identify the follicular Tr1–FCRL4⁺ MBC axis as a central immunoregulatory circuit in pediatric tonsillar hypertrophy. This pathway maintains a hyporesponsive yet persistent FCRL4⁺ MBC pool, reshaping the follicle through reciprocal signaling, transcriptional checkpoints, and spatial organization. By defining this tolerance-based circuit our study connects follicular immunoregulation with pediatric airway pathology, establishing a translational rationale for immunomodulatory approaches that may ultimately reduce adenotonsillectomy dependence.

## Methods

### Study design and patient cohort

This study was designed as a prospective, multi-platform translational investigation to elucidate immune mechanisms underlying pediatric tonsillar hypertrophy (TH). Seventy-seven children aged 3– 17 years undergoing tonsillectomy at Seoul National University Children’s Hospital were enrolled after obtaining informed consent from legal guardians. All procedures were approved by the Institutional Review Board of Seoul National University Hospital (IRB No. H-2209-050-1357). Participants were categorized into small (Brodsky grade < 3; n = 40) and big (grade ≥ 3; n = 37) tonsil groups, with demographic and clinical characteristics balanced between groups (Table 1).

A tiered discovery–validation framework integrating single-cell and spatial multi-omic analyses was employed to ensure reproducibility across analytic layers. The Discovery-Main cohort (n = 16; 8 small and 8 big) underwent comprehensive profiling, including single-cell RNA sequencing, GeoMx Digital Spatial Profiling, PhenoCycler-Fusion multiplex imaging, immunohistochemistry for CD20 and FCRL4, and confirmatory flow cytometry. The Discovery-Extended cohort (n = 29; 15 small and 14 big) provided additional analyses through immunohistochemistry for CD20, flow cytometry and MACSima spatial proteomics. The independent Validation cohort (n = 32; 17 small and 15 big) was analyzed exclusively by single-cell RNA sequencing using identical pipelines and integrated via label transfer (Fig. 1A; Supplementary Table 1). All analyses were performed blinded to clinical metadata until completion of cell-type annotation.

### Tonsil sample processing

#### Tissue dissociation and mononuclear cell isolation

Pediatric tonsil tissue was washed with phosphate-buffered saline (PBS), cut into small pieces on a 60-mm Petri dish, and mechanically minced with sterile scissors. Approximately three-eighths of each tonsil was then subjected to enzymatic digestion in Liberase TL (2 mg/mL; Roche, Cat. No. 05401020001) diluted in PBS at 37 °C with gentle agitation. The digested suspension was gently triturated and passed through a 70-µm strainer into a conical tube containing serum-containing buffer. Cells were pelleted by centrifugation, resuspended in PBS, and isolated using Ficoll-Paque density-gradient centrifugation (Cytiva, Cat. No. 17144003). Mononuclear cells were washed, pelleted, and resuspended in Cell Banker 1 (ZENOAQ, Cat. No. BLC-1) before counting, aliquoting, and cryopreservation at −80 °C followed by long-term storage in liquid nitrogen.

#### FFPE tissue preparation and tissue microarray construction

To support spatial transcriptomic and immunohistochemical analyses, tissue microarrays (TMAs) were generated by extracting a single 2-mm core from a representative region of formalin-fixed, paraffin-embedded (FFPE) tonsil tissue for each patient. This approach enabled standardized and region-specific comparison of tissue architecture and molecular features across samples.

### Single-cell RNA sequencing

#### Library preparation and sequencing

Single-cell RNA sequencing was performed using the 10x Genomics Chromium 5’ v2 HT protocol (10X Genomics, Cat. No. CG000424). Single-cell suspensions from 48 pediatric tonsil samples were filtered through a 40 µm cell strainer. For cell labeling, the cell number of each sample was adjusted to 1,000,000 cells in RPMI-1640 with 10% FBS. To block Fc receptors and reduce non-specific binding, the cells were pre-treated with Human TruStain FcX (BioLegend, Cat. No. 422301). Each cell was labeled with TotalSeq anti-human Hashtag Antibody (Biolegend) for multiplexing. Subsequently, cells were loaded onto a Chromium X using the 10x Genomics 5’ v2 HT Chip. Cells were encapsulated in droplets containing gel beads and partitioning oil. Reverse transcription was performed within the droplets to synthesize complementary DNA (cDNA), followed by cDNA amplification according to the 10x Genomics protocol. Gene Expression (GEX) libraries were then prepared using the 10x Genomics 5’ v2 HT reagents, which involved cDNA fragmentation, amplification, and the addition of sequencing adapters. Additionally, BCR libraries were constructed using the 10x Genomics BCR enrichment kits, following the manufacturer’s instructions for enrichment and preparation of BCR sequences from the cDNA. The prepared libraries were evaluated for quality using an Agilent Bioanalyzer. Libraries were assessed for quality using an Agilent Bioanalyzer. Sequencing was performed on an Illumina platform, targeting sufficient read depth to ensure comprehensive profiling and accurate HashTag label identification.

#### Data processing and quality control

For 10X sequencing data, the Cell Ranger pipeline (v 7.1.0) provided by 10X Genomics was applied to align reads and produce the gene-cell unique molecular identifier (UMI) matrix. Seurat (v5.1.0) was used to read the scRNA-seq expression matrix^81^. The raw reads were demultiplexed using the hashtag oligo (HTO) libraries and “HTODemux” function with default parameters from Seurat. Features detected in fewer than three cells were excluded, and cells that expressed below 200 genes or over 10% mitochondrial genes were removed. scDblFinder package (version 1.18) was used for the identification and removal of doublets in the sample^82^. Gene expression levels were normalized with “NormalizedData” function. The Harmony algorithm was performed to remove the batch effect^83^. The top 2000 highly variable genes were identified via the FindVariableFeatures function. Principal component analysis (PCA) was performed using the top 2000 highly variable genes with RunPCA function, and the top 20 principal components were used for Uniform Manifold Approximation and Projection for Dimension Reduction (UMAP). The FindNeighbors and FindClusters function was further performed for the unsupervised clustering. Cell types were annotated based on the canonical marker gene expression, found using the FindAllMarkers function with the parameter “min.pct = 0.25”. First, broad cell type was defined and each cell type was further sub-clustered with the same procedures.

### Single-cell RNA sequencing data analysis

#### B cell receptor (BCR) sequencing analysis

The BCR sequence data was processed using the Immcantation (immcantation.org) pipeline with default parameters. Briefly, filtered V(D)J contigs for BCR sequence data were obtained by Cell Ranger. IgBLAST (www.ncbi.nlm.nih.gov/igblast/) database was used to assign V(D)J gene annotations with Change-O package. Nonproductive heavy and light chain BCR sequences were removed. Cells with multiple heavy chains were removed. BCRs were annotated by B cell subsets based on matching their single-cell barcodes to the gene expression information. Somatic hypermutation (SHM) counts were calculated using shazam (v1.2.0) based on different nucleotide bases between sequence alignments against germline-d-mask-alignments.

#### Differentially expression and enrichment analysis

Differentially expressed genes (DEGs) under different conditions were calculated among the genes using the “FindMarkers” functions from Seurat with the wilcoxon rank sum test. Genes with an adjusted p value < 0.05 and a |log2 fold change| > 1 were considered as DEGs. Gene Ontology (GO) enrichment analysis was performed on the DEGs using the R package ClusterProfiler to identify enriched functional pathways between the groups.

#### Signature gene set scoring

The signature gene sets for scoring were identified by using FindAllMarkers function with the parameter “min.pct = 0.25” and selecting genes with adjusted p value < 0.05 and log2 fold change > 1 (Follicular helper T cell and Follicular Tr1 cell) or 2 (FCRL4^+^ Memory B cell) for each cell cluster. Visualization for the score in UMAP plot was performed by using “AddModuleScore” from Seurat.

#### Cell-cell interaction analysis

To analyze cell-cell interactions between clusters, we employed the CellPhoneDB database with normalized count data as an input file. The significant ligand-receptor pairs were filtered with a P value of less than 0.05^84^.

#### Trajectory analysis

For trajectory analysis of the T/NK population, the Velocyto package (v0.17.17) was used to annotate reads as spliced or unspliced based on their genomic and transcriptomic alignments^85^. RNA velocity was estimated using scVelo (v0.3.2), which applies a stochastic model to infer splicing dynamics^86^. For the B cell population, trajectory analysis was performed using the partition-based graph abstraction (PAGA) approach implemented in the Scanpy package (v1.9.8)^87^.

#### Validation cohort analysis

The single-cell sequencing data for the validation cohort was processed using the same procedure as that applied to the discovery cohort. The SingleR package (v2.6.0) was used to identify cell populations and infer cell types, using the scRNA-seq data from the discovery cohort as the reference dataset^88^.

### Spatial Transcriptomic Profiling

#### Tissue preparation and GeoMx DSP assay

Spatial transcriptomic profiling was performed using the GeoMx Digital Spatial Profiler (DSP) (NanoString)^89^. Four-micrometer FFPE tonsil sections were mounted on charged slides (Leica BOND Plus), deparaffinized, and rehydrated according to the manufacturer’s protocol. Slides were incubated with RNA detection probes containing UV-photocleavable oligonucleotide barcodes, followed by fluorescent staining with SYTO 13 (NanoString, Cat. No. 121300303), Pan-cytokeratin (PanCK; Novus, Cat. No. NBP2-33200AF532), CD20 (Novus, Cat. No. NBP2-47840DL594), and CD3 (Novus, Cat. No. NBP2-54392AF647). After whole-slide imaging, one region of interest (ROI) per patient was selected and segmented into CD20-positive and CD20-negative compartments, yielding a total of 32 segments. Oligonucleotide tags were released by UV illumination, collected into a 96-well plate, indexed during library preparation, and pooled for next-generation sequencing–based digital quantification.

#### Data processing, normalization, and analysis

Sequencing data were processed using the GeoMx DSP Analysis Suite (v3.1.2.12) to generate digital count matrices. Regions and genes that did not meet default quality-control criteria were excluded, and count data were normalized using the third-quartile (Q3) method. Differentially expressed genes (DEGs) were identified using unpaired t-tests with Benjamini–Hochberg correction for multiple comparisons. Gene Ontology (GO) enrichment analysis was performed using the clusterProfiler package in R. Normalized data were further analyzed using Gene Set Enrichment Analysis (GSEA) and single-sample GSEA (ssGSEA) with cell-type signatures derived from the single-cell RNA-seq dataset.

### Tissue preparation and Xenium assay

For single-cell–resolved spatial transcriptomic analysis, tonsil tissues were profiled using the Xenium In Situ platform (10x Genomics) according to the manufacturer’s protocol. Briefly, 5-μm formalin-fixed, paraffin-embedded (FFPE) tonsil sections were mounted on Xenium slides, deparaffinized, and subjected to target retrieval. Tissue sections were hybridized with the Xenium Human Immuno-Oncology (hIO) Profiling Panel (380 genes), supplemented with a custom 50-gene panel, followed by probe ligation, rolling-circle amplification, and iterative fluorescence imaging on the Xenium Analyzer. Cell segmentation and transcript assignment were performed using the Xenium Onboard Analysis pipeline.

### Xenium data processing and analysis

Xenium data were processed using stLearn (v1.1.1) and Scanpy (v1.10.4). Genes and cells with fewer than 10 total counts were filtered out, and the count matrix was normalized by total counts. The normalized expression matrix was transformed using a square-root transformation, followed by principal component analysis. A neighborhood graph was constructed using the top 50 principal components, and cells were clustered using the Leiden algorithm. Cell types were annotated based on canonical marker gene expression patterns. To further support the annotation, cell type signature gene sets were derived from the single-cell RNA-seq data using genes overlapping with the Xenium gene panel, and signature scores were calculated in the Xenium dataset. To assess spatial interactions between cell types, neighborhood enrichment analysis was performed using Squidpy (v1.7.0) based on cell type annotations. Cell localization and spatial distribution patterns were visualized using Xenium Explorer.

### Multiplexed spatial protein imaging

#### PhenoCycler-Fusion multiplex imaging

FFPE tonsil tissue blocks from patients with matched scRNA-seq data were analyzed using the PhenoCycler-Fusion platform (formerly CODEX; Akoya Biosciences)^90^. Images were analyzed using QuPath (v0.5.1)^91^. For each patient, up to three follicles and their surrounding regions were selected as ROIs. CD20⁺ B cells were quantified based on co-localized DAPI and CD20 expression within each ROI.

#### MACSima cyclic immunofluorescence imaging

Multiplexed spatial protein profiling was performed using the MACSima Imaging System (Miltenyi Biotec), an automated cyclic immunofluorescence platform^92^. FFPE tonsil sections were processed according to the manufacturer’s protocol, including antigen retrieval. A sequential panel of fluorophore-conjugated antibodies was applied through iterative cycles of staining, image acquisition, and fluorophore inactivation. High-resolution images were collected across multiple spectral channels and analyzed using MACS iQ View software for cell segmentation, marker quantification, and spatial profiling. ROIs were defined based on tonsillar follicular architecture, including germinal centers, mantle zones, and adjacent interfollicular regions, to characterize the local immune microenvironment and spatial distribution of immune cell subsets.

### Immunohistochemistry

#### CD20 staining

Unstained 4 μm-thick sections were prepared from FFPE tonsil tissue blocks for immunohistochemical analysis. For CD20 staining, sections were incubated with a mouse monoclonal anti-CD20 antibody (clone L26; Dako) and processed on the Bond-Max Autostainer (Leica Microsystems) according to the manufacturer’s instructions. Whole-slide images were acquired using the Aperio GT450 scanner (Leica Microsystems) at 40× magnification, and follicle size measurements were manually assessed by a pathologist (J.K.).

#### FCRL4 staining

For FCRL4 staining, sections were processed on the BenchMark ULTRA system (Roche Diagnostics) using a rabbit monoclonal anti-FCRL4 antibody [EPR21961] (Abcam, Cat. No. ab239076) and the Optiview DAB detection system, following the manufacturer’s protocol. FCRL4⁺ cells were quantified using QuPath (v0.5.1).

### Flow Cytometry Analysis

Single-cell suspensions from tonsil tissue were prepared and stained with fluorophore-conjugated monoclonal antibodies in DPBS containing 2% FBS. To exclude dead cells, samples were first incubated with Fixable Viability Stain 700 (BD Biosciences) at a 1:1000 dilution in DPBS for 15 minutes at 4°C, followed by blocking with Human TruStain FcX (BioLegend) to prevent nonspecific Fc receptor binding. Surface staining was performed for 30 minutes at 4°C in the dark. For intracellular Foxp3 detection, cells were fixed and permeabilized using the Foxp3/Transcription Factor Staining Buffer Set (eBioscience), according to the manufacturer’s instructions. The following antibodies were used: BUV496 anti-human CD4 (Clone SK-3), APC-H7 anti-human CD8 (HIT8a), PE anti-human Foxp3 (236A/E7), BUV737 anti-human CD25 (2A3), BV786 anti-human CD127 (HIL7R-M21), BV421 anti-human CXCR5 (FR8B2), BV711 anti-human PD-1 (EH122.1), Alexa Fluor 488 anti-human BCL6 (K112-91) (BD Biosciences); BV605 anti-human CD3 (OKT3) (BioLegend); and PE-Cy7 anti-human ICOS (ISA-3) (eBioscience). Samples were acquired using a BD Symphony A3 flow cytometer (BD Biosciences), and data were analyzed using FlowJo software (version 10.9.0, BD Biosciences).

## Statistical analysis

All statistical analyses were performed using R version 4.4.0 and GraphPad Prism 9 (GraphPad Software). The Spearman rank correlation was used to evaluate associations between variables, while group comparisons were performed using the Mann–Whitney U test for unpaired data. P values < 0.05 were considered statistically significant.

## Data availability

All sequencing data are being uploaded to GEO. The accession number will be provided during review and made public upon publication.

## Code availability

Relevant codes used to perform the analysis and reproduce the figures are available on GitHub: https://github.com/AkiGeonWooPark/Integrated-scRNA-and-GeoMx-analyis-for-Tonsillar-Hypertrophy

## Table/Figure Legend

Supplementary Table 1. Clinical metadata and experimental platforms for each pediatric tonsil sample.

Supplementary Table 2. Differentially expressed genes (DEGs) of FCRL4⁺ memory B cells (MBCs) across tonsil subgroups.

Supplementary Table 3. Gene Ontology (GO) enrichment analysis of DEGs identified in FCRL4⁺ MBCs.

## Acknowledgement

NA

## Author Contributions

J.Y.B. processed samples, performed experiments, analyzed data, and wrote the manuscript. G.W.P. performed data analysis and contributed to manuscript writing. S.J. managed clinical data and contributed to manuscript writing. J.K. performed histopathological analyses. Y.M., H.S., H.Y.S., C.K., and T.G. processed samples and performed experiments. M.K. reviewed and revised the manuscript. D.H.H. conceived and supervised the study. H.J.K. supervised the study, acquired funding, and provided overall project guidance. All authors reviewed and approved the final version of the manuscript.

## Additional information

Supplementary information

Supplementary table

---

**Supplementary Figure 1.**
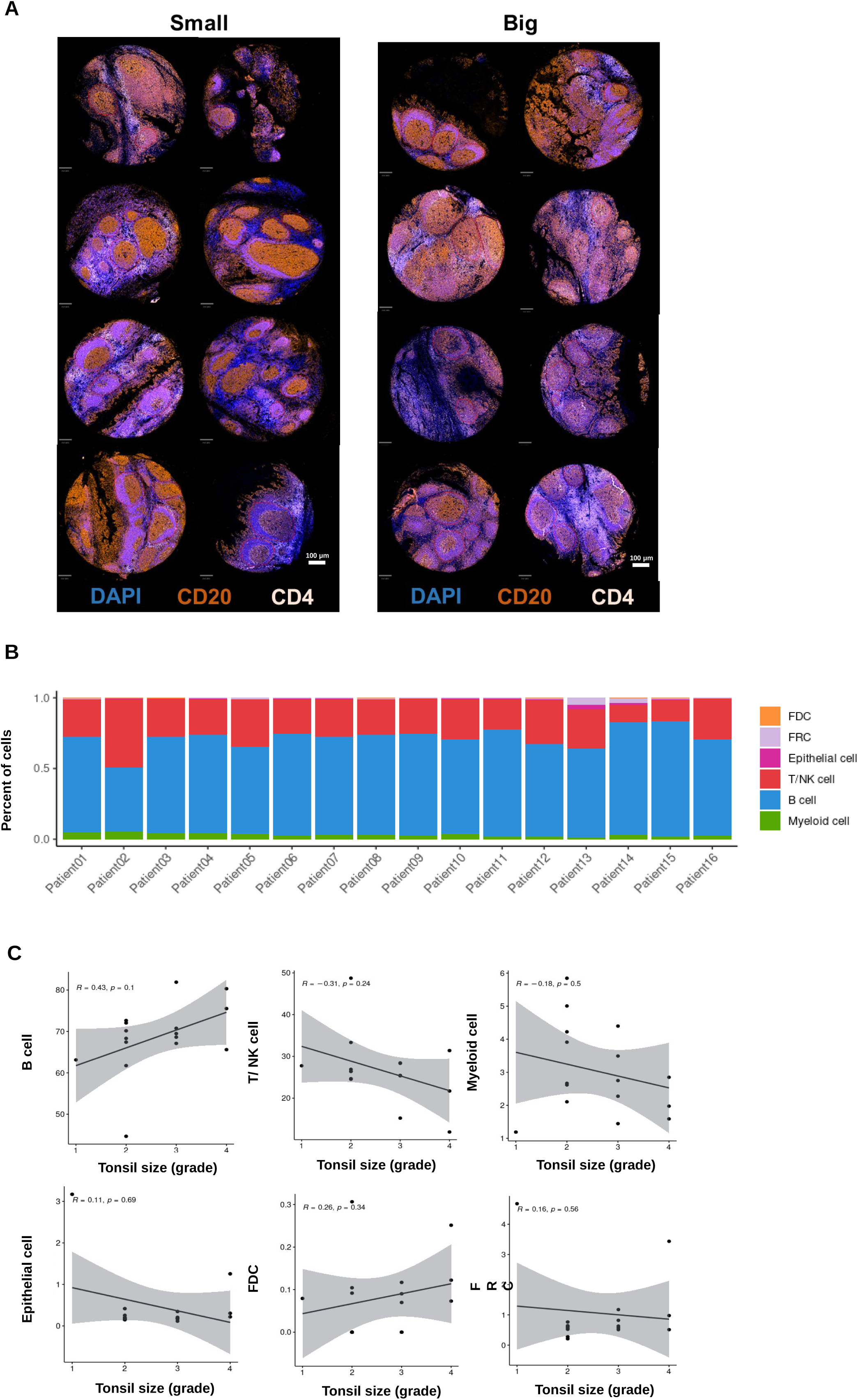
Spatial imaging and cell-type composition of pediatric tonsils. (A) Multiplex immunofluorescence image of pediatric tonsil tissue from the Discovery-Main cohort (small, n = 8; big, n = 8), acquired using the Akoya platform and showing DAPI (nuclei), CD20 (B cells), and CD4 (T cells). (scale bars, 250 μm). (B) Proportions of major immune cell types identified by scRNA-seq across the Discovery-Main cohort (n = 16). Each bar represents one patient. (C) Correlation between pre-operative right tonsil size and the relative frequencies of immune cell types across the Discovery-Main cohort (small, n = 8; big, n = 8). Correlation coefficients (R) and p-values are indicated in each panel. No statistically significant associations were observed.

**Supplementary Figure 2.**
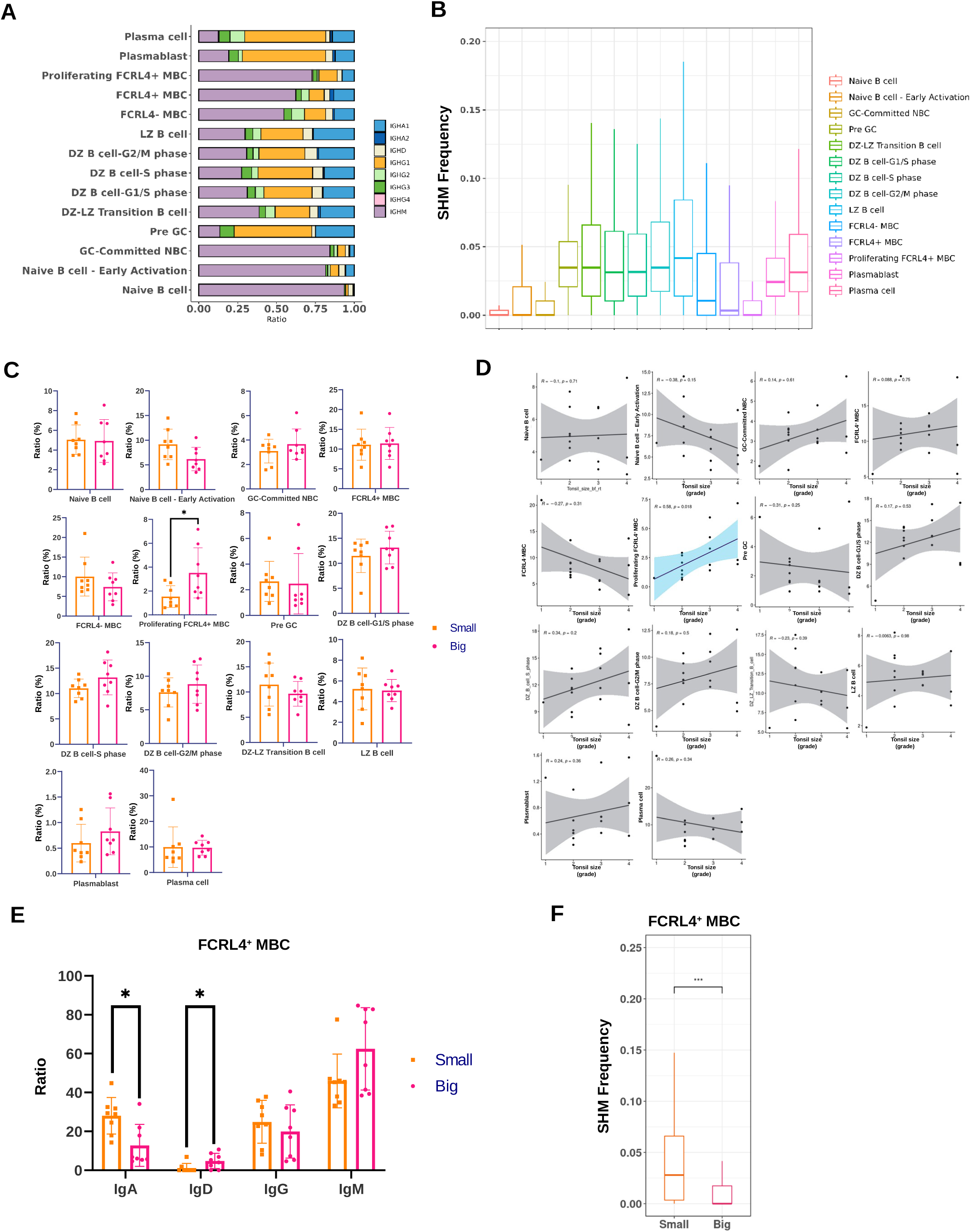
Differential features of FCRL4⁺ MBCs in hypertrophic tonsils. (A) Isotype composition across B cell subsets identified by scRNA-seq in the Discovery-Main cohort (n = 16). (B) SHM frequency across B cell subsets in the Discovery-Main cohort (n = 16). Box plots indicate the distribution of mutation frequencies per subset. (C) Proportions of B cell subsets in tonsils from the Discovery-Main cohort (small, n = 8; big, n = 8). Each dot represents one patient. Bars indicate mean ± SD. (D) Correlation between pre-operative right tonsil size and the frequencies of B cell subsets in the Discovery-Main cohort (small, n = 8; big, n = 8). Each dot represents one patient. (E) Isotype switching frequency in FCRL4⁺ MBCs from tonsils in the Discovery-Main cohort (small, n = 8; big, n = 8). Each dot represents one patient. Bars indicate mean ± SD. (F) SHM frequency in FCRL4⁺ MBCs from tonsils in the Discovery-Main cohort (small, n = 8; big, n = 8). Box plots indicate the distribution of mutation frequencies per group. Statistical significance was assessed using Mann–Whitney test in (C), (E), and (F) (*p < 0.05, ***p < 0.001) and Spearman correlation in (D). SHM, somatic hypermutation; MBC, memory B cell.

**Supplementary Figure 3.**
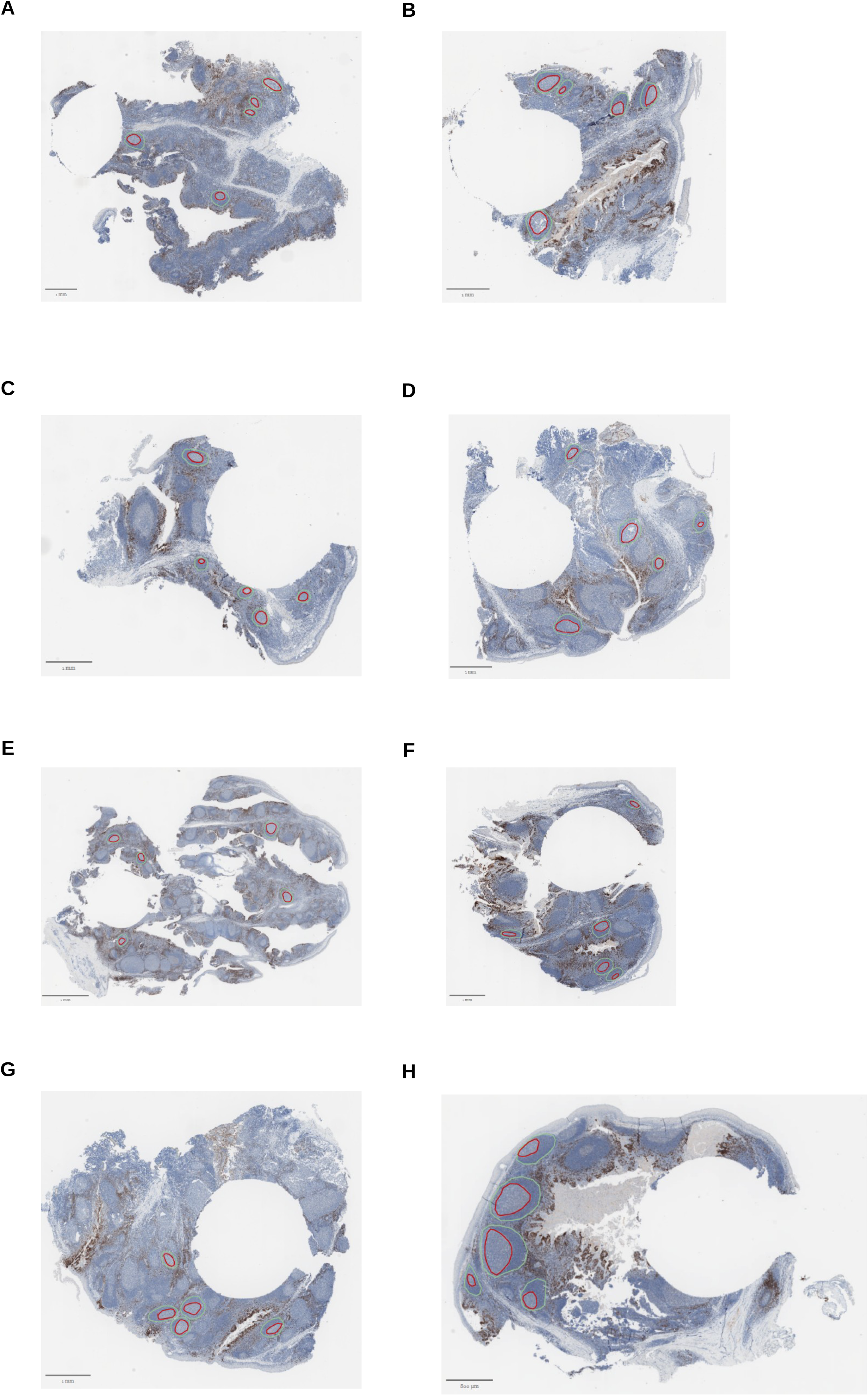
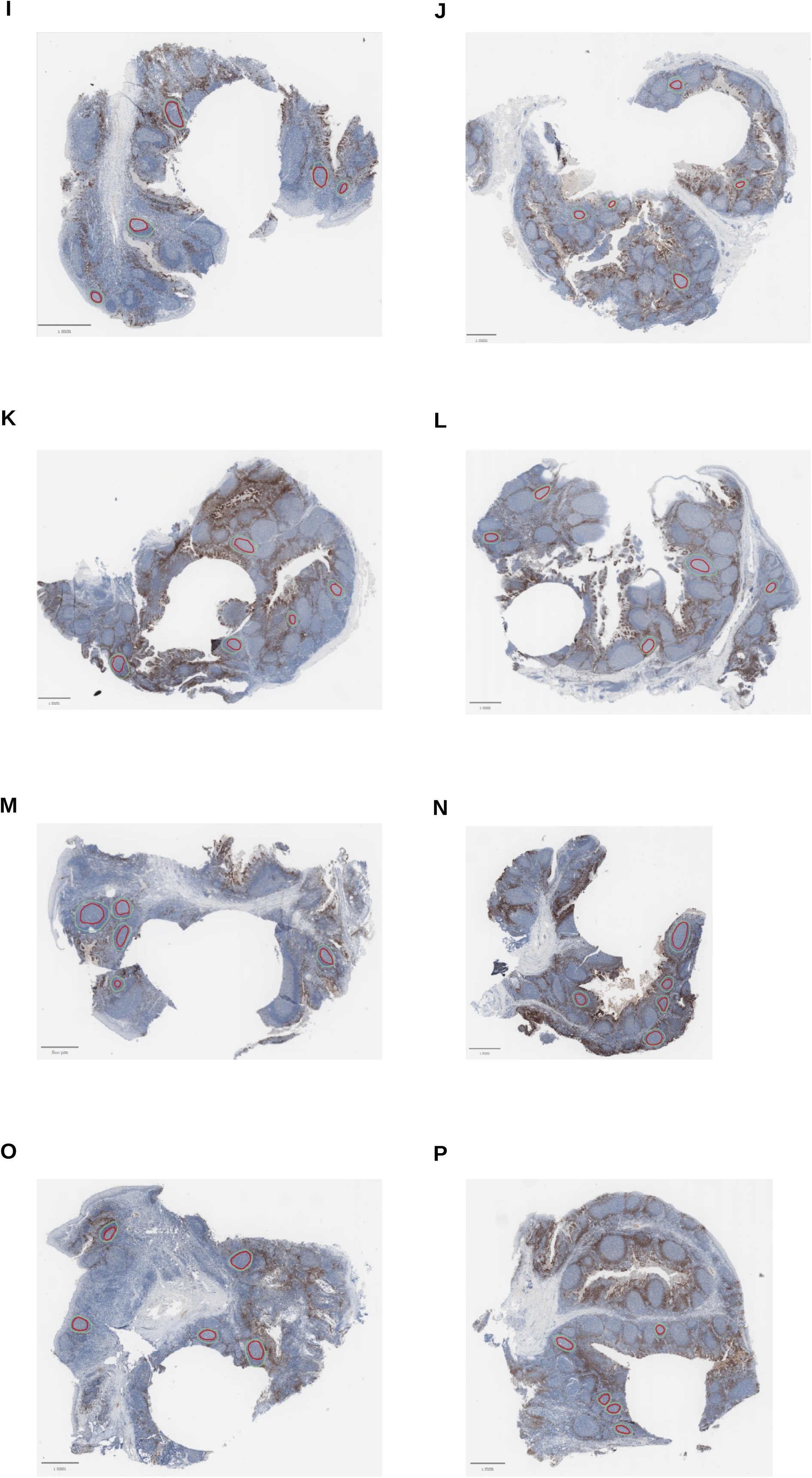
FCRL4 immunohistochemistry in tonsils from the Discovery-Main cohort. (A–H) Whole-slide immunohistochemistry images of tonsillar sections from small tonsils (n = 8). (I–P) Whole-slide immunohistochemistry images of tonsillar sections from big tonsils (n = 8). Five follicles per sample were annotated, with germinal centers outlined in red and follicular boundaries in green. Scale bars are indicated in each panel and may vary across samples.

**Supplementary Figure 4.**
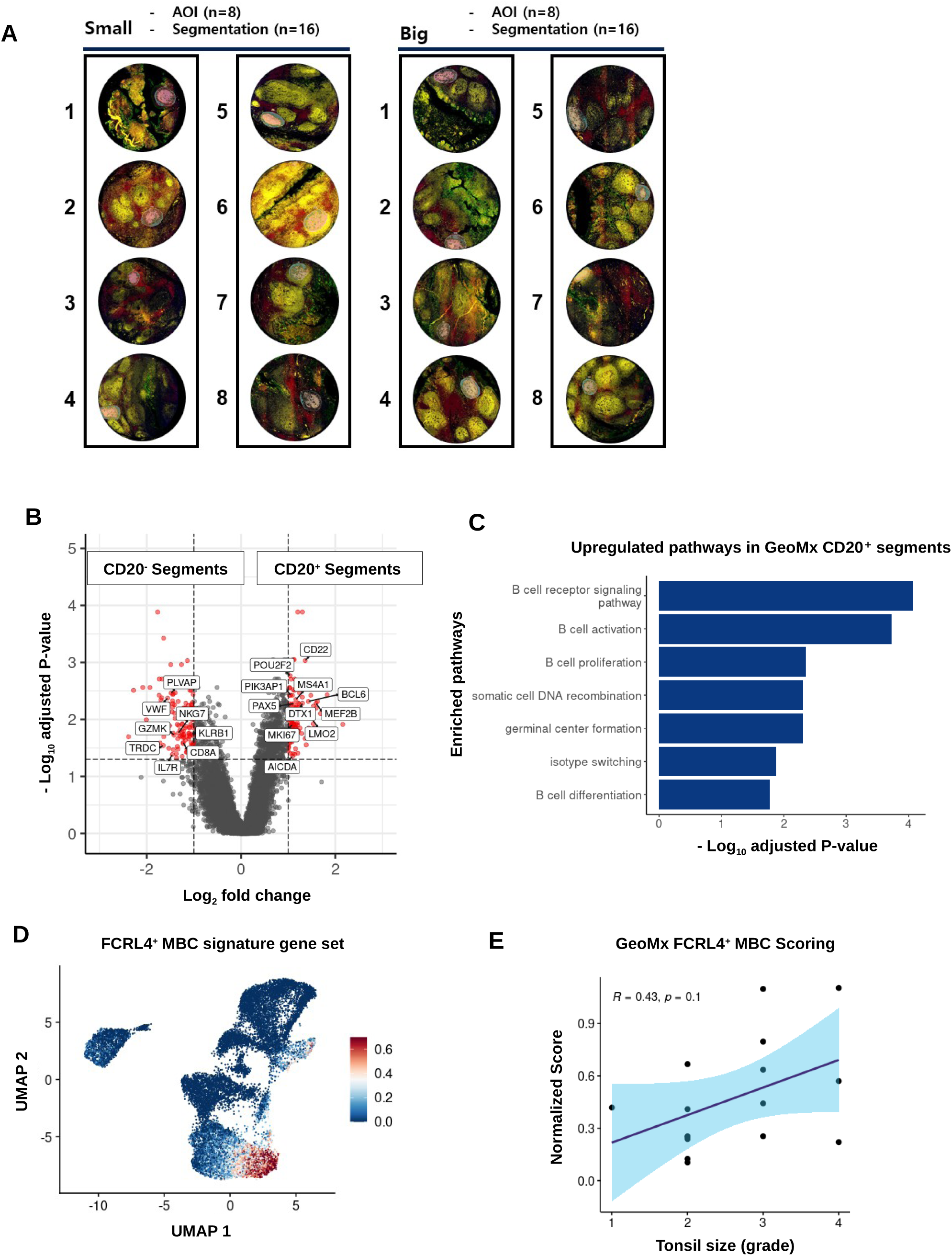
Spatial validation of the FCRL4⁺ MBC signature in tonsils by GeoMx DSP. (A) GeoMx DSP image of pediatric tonsil tissue from the Discovery-Main cohort (small, n = 8; big, n = 8), stained with SYTO13 (nuclei), Pan-CK (epithelial cells), CD20 (B cells), and CD3 (T cells), segmented into CD20⁺ and CD20⁻ compartments (scale bars, 200 μm). (B) Volcano plot of DEGs between CD20⁺ and CD20⁻ segments, with genes upregulated in CD20⁺ segments highlighted. (C) Pathway enrichment analysis of genes upregulated in CD20⁺ segments. (D) UMAP of scRNA-seq data showing expression of the FCRL4⁺ MBC signature gene set across B cell subsets in the Discovery-Main cohort (small, n = 8; big, n = 8). (E) Correlation between GeoMx ssGSEA scores for the FCRL4⁺ MBC signature in CD20⁺ segments and pre-operative right tonsil size in the Discovery-Main cohort (small, n = 8; big, n = 8). Each dot represents one patient. Statistical significance was assessed using Spearman correlation in (E). DSP, digital spatial profiling; DEGs, differentially expressed genes; UMAP, Uniform Manifold Approximation and Projection; ssGSEA, single-sample Gene Set Enrichment Analysis; MBC, memory B cell.

**Supplementary Figure 5.**
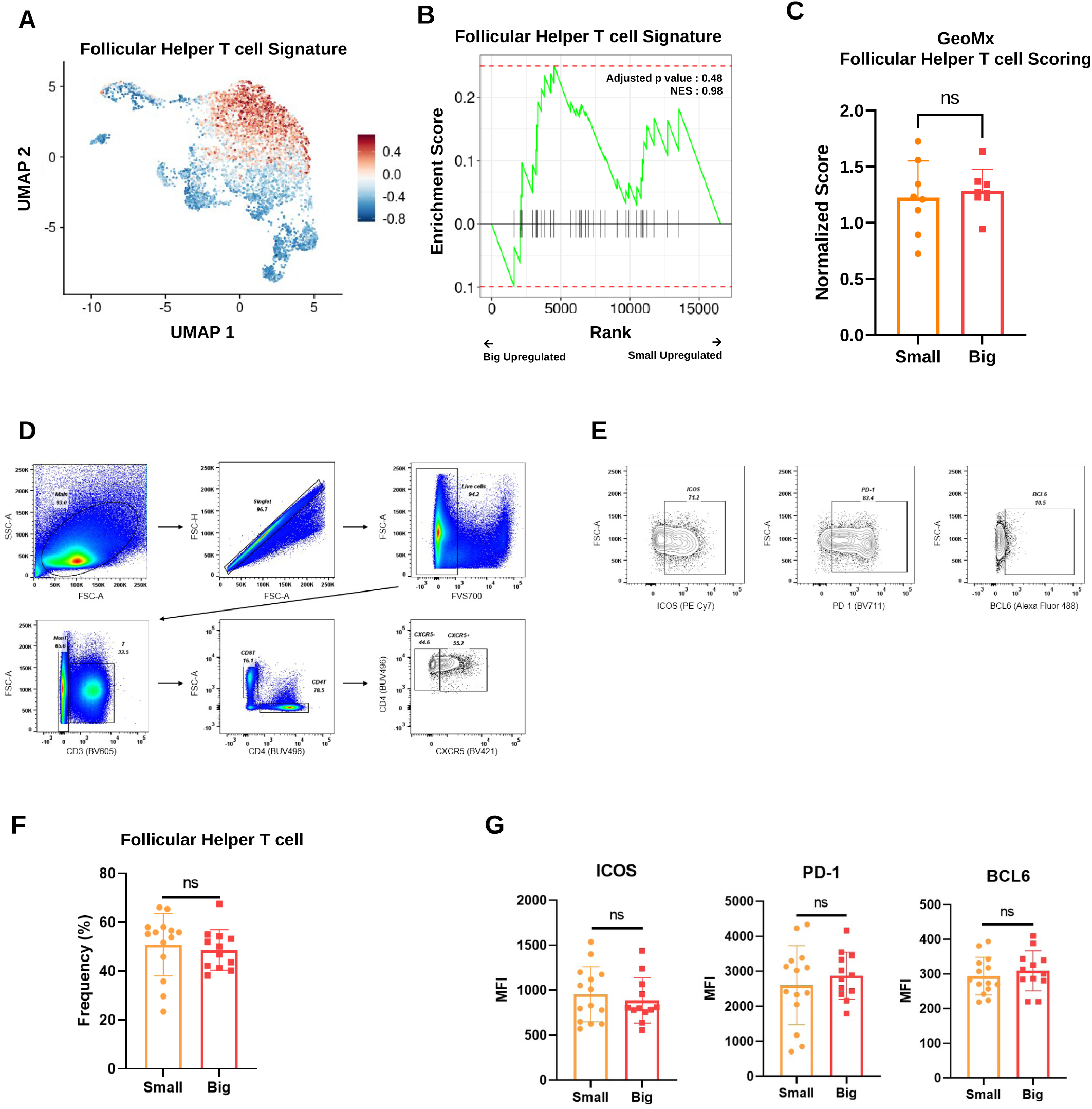
Spatial and transcriptomic evaluation of Tfh signatures. (A) UMAP of scRNA-seq data showing expression of the Tfh signature gene set across T/NK subsets in the Discovery-Main cohort (small, n = 8; big, n = 8). (B) GSEA of the Tfh signature in CD20⁻ segments from tonsils in the Discovery-Main cohort (small, n = 8; big, n = 8). (C) ssGSEA scores for the Tfh signature in CD20⁻ segments from tonsils in the Discovery-Main cohort (small, n = 8; big, n = 8). Each dot represents one patient. Bars indicate mean ± SD. (D) Flow cytometry gating strategy for Tfh cells. Live lymphocytes were gated by FSC/SSC and singlets by FSC-A/FSC-H, followed by exclusion of dead cells (FVS700⁺). CD3⁺ T cells were selected, and CD4⁺ cells were identified within this population. Tfh cells were then defined as CXCR5⁺CD4⁺ T cells. (E) Representative expression of ICOS, PD-1, and BCL6 in gated Tfh cells. (F) Flow cytometric quantification of Tfh cell frequencies in tonsils (small, n = 14; big, n = 12). Each dot represents one patient. Bars indicate mean ± SD. (G) Flow cytometric comparison of ICOS, PD-1, and BCL6 expression in Tfh cells (small, n = 14; big, n = 12). Each dot represents one patient. Bars indicate mean ± SD. Statistical significance was assessed using GSEA in (B) and Mann–Whitney test in (C), (F), and (G); no significant differences were observed. UMAP, Uniform Manifold Approximation and Projection; Tfh, T follicular helper cell; GSEA, Gene Set Enrichment Analysis; ssGSEA, single-sample Gene Set Enrichment Analysis.

**Supplementary Figure 6.**
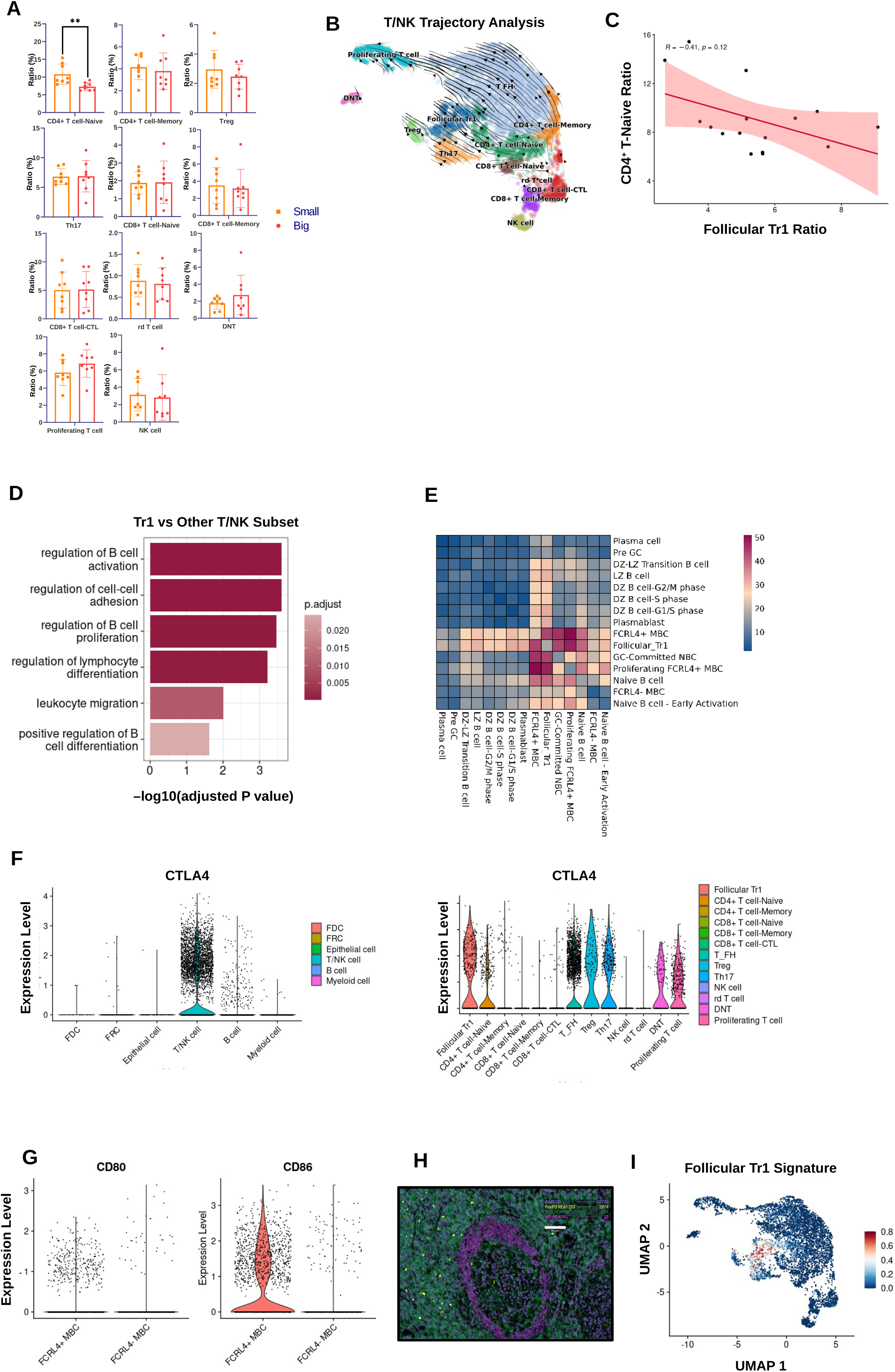
Transcriptional, interaction, and spatial features of follicular Tr1 cells in hypertrophic tonsils. (A) Frequencies of T cell subsets from tonsils in the Discovery-Main cohort (small, n = 8; big, n = 8). Each dot represents one patient. Bars indicate mean ± SD. (B) RNA velocity analysis of tonsillar T/NK cells showing transcriptional trajectories toward follicular Tr1 cells. (C) Correlation between follicular Tr1 and naive CD4⁺ T cell frequencies in the Discovery-Main cohort (small, n = 8; big, n = 8). Each dot represents one patient. (D) Pathway enrichment analysis of DEGs in follicular Tr1 cells compared to other T/NK subsets. (E) Cell–cell interaction analysis between follicular Tr1 cells and B cell subsets. (F) Violin plot analysis of CTLA-4 expression across major immune lineages (left) and within T cell subsets (right) in tonsils from the Discovery-Main cohort (small, n = 8; big, n = 8). (G) Violin plot analysis of CD80 and CD86 expression in FCRL4⁺ versus FCRL4⁻ memory B cells from tonsils in the Discovery-Main cohort (small, n = 8; big, n = 8). (H) Multiplex immunofluorescence image (MACSima) of tonsillar follicles stained for DAPI (purple), FOXP3 (yellow), CD4 (green) and IgD (magenta) (scale bars, 200 μm). (I) UMAP visualization of scRNA-seq data from the Discovery-Main cohort, colored by follicular Tr1 signature score. Statistical significance was assessed using Mann–Whitney test in (A) (**p < 0.01) and Spearman correlation in (C). Tr1, type 1 regulatory T cell; DEGs, differentially expressed genes; UMAP, Uniform Manifold Approximation and Projection.

**Supplementary Figure 7.**
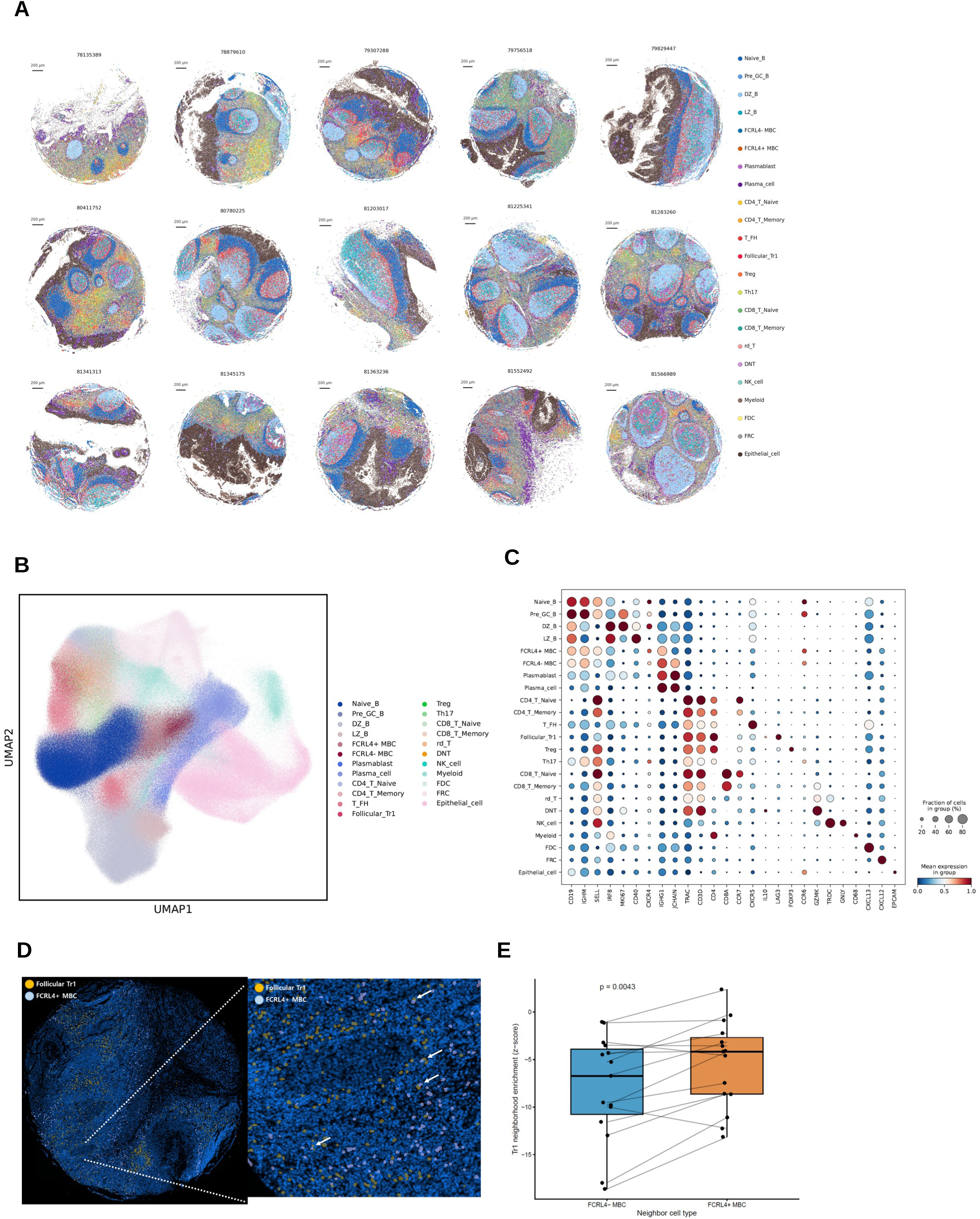
Spatial association between follicular Tr1 cells and FCRL4⁺ memory B cells in human tonsils. (A) Spatial distribution of annotated cell populations across 15 tonsillar tissue samples profiled by Xenium. (B) UMAP representation of Xenium-resolved cells colored by cell-type annotation. (C) Dot plot showing the expression of canonical marker genes used to define the 23 annotated cell populations. Dot size indicates the fraction of cells expressing each gene, and color indicates mean expression within each cell population. (D) Representative spatial visualization showing the localization of follicular Tr1 cells and FCRL4⁺ MBCs within tonsillar tissue, with an enlarged view highlighting their close spatial apposition. (E) Neighborhood enrichment analysis comparing the association of follicular Tr1 cells with FCRL4⁻ versus FCRL4⁺ MBCs. Lines connect paired measurements from the same tissue sample.

**Supplementary Figure 8.**
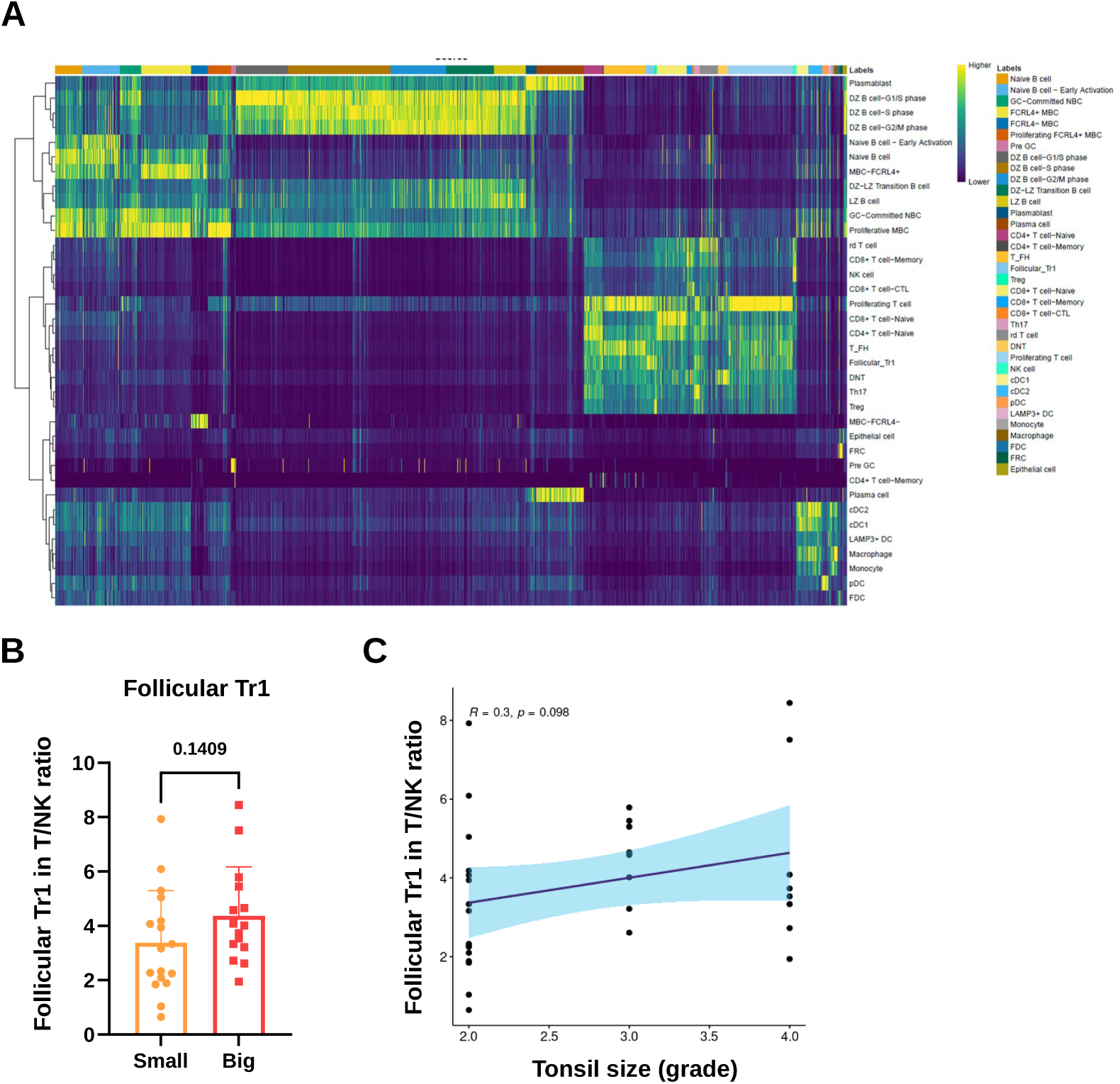
Validation of follicular Tr1 cell frequency in an independent cohort. (A) Heatmap of SingleR-based automated cell type annotation in the Validation cohort (n = 32) using the Discovery-Main cohort dataset (n = 16) as reference. (B) Proportion of follicular Tr1 cells among T/NK cells in tonsils from the Validation cohort (small, n = 17; big, n = 15). Each dot represents one patient. Bars indicate mean ± SD. (C) Correlation between follicular Tr1 frequency and pre-operative right tonsil size in the Validation cohort (small, n = 17; big, n = 15). Statistical significance was assessed using Mann–Whitney test in (B) and Spearman correlation in (C); no significant differences were observed. Tr1, type 1 regulatory T cell.

